# *In vivo* intra-uterine delivery of TAT-fused Cre recombinase and CRISPR/Cas9 editing unveil histopathology of Pten/p53-deficient endometrial cancers

**DOI:** 10.1101/2023.06.08.544224

**Authors:** Raúl Navaridas, Maria Vidal-Sabanés, Anna Ruiz-Mitjana, Gisela Altés, Aida Perramon-Güell, Andree Yeramian, Joaquim Egea, Mario Encinas, Sonia Gatius, Xavier Matias-Guiu, Xavier Dolcet

**Affiliations:** Developmental and Oncogenic Signalling Group. Departament de Ciències Mèdiques Bàsiques and Departament de Medicina Experimental. Universitat de Lleida. Institut de Recerca Biomèdica de Lleida, IRBLleida. Lleida, Spain; Oncologic Pathology Group. Departament de Ciències Mèdiques Bàsiques, Universitat de Lleida. Institut de Recerca Biomèdica de Lleida, IRBLleida. Lleida, CIBERONC, Spain

**Keywords:** cell penetrating peptides, HIV1-TAT, Cre-recombinase, mouse model of cancer, Pten, p53, CRISPR/Cas9, endometrial cancer, uterine carcinosarcoma

## Abstract

Pten and p53 are two of the most frequently mutated tumor suppressor genes in endometrial cancer. However, the functional consequences and histopathological manifestation of concomitant p53 and Pten loss of function alterations in the development of endometrial cancer is still controversial. Here, we demonstrate that simultaneous Pten and p53 deletion is sufficient to cause epithelial to mesenchymal transition phenotype in endometrial organoids. By a novel TAT-fused Cre intravaginal delivery method, we achieved local ablation of both p53 and Pten specifically in the uterus. These mice developed high-grade endometrial carcinomas and a high percentage of uterine carcinosarcomas resembling those found in humans. To further demonstrate that carcinosarcomas arise from epithelium, double Pten/p53 deficient epithelial cells were mixed with wild type stromal and myometrial cells and subcutaneously transplanted to Scid mice. All xenotransplants resulted in the development of uterine carcinosarcomas displaying high nuclear pleomorphism and metastatic potential. Accordingly, in vivo CRISPR/Cas9 disruption of Pten and p53 also triggered the development of metastatic carcinosarcomas. Our results unfadingly demonstrate that simultaneous deletion of p53 and Pten in endometrial epithelial cells is enough to trigger epithelial to mesenchymal transition that is consistently translated to the formation of uterine carcinosarcomas in vivo.

## INTRODUCTION

Endometrial cancer (EC) is the most frequent type of cancer in the female reproductive tract [1] [2]. EC can be classified from pathological, histological or molecular point of view[3]. Traditionally, EC has been classified into two main groups on basis of clinical and pathological features: Type I or endometroid endometrial carcinoma (ECC) and Type II or non-endometroid endometrial carcinomas (NECC)[4]. Histologically, the main appearances of EC are endometroid carcinomas (including its variants), clear cell carcinomas, serous carcinomas or uterine carcinosarcomas (UCS)[1,3]. The latter ones are biphasic malignant tumors containing carcinomatous (epithelial) and sarcomatous (mesenchymal) compartments[5]. Although molecular profiling of EC supports that mesenchymal cells of the sarcomatous compartment arise from epithelial-to-mesenchymal trans-differentiation (EMT) of epithelial cells, there is no current functional validation for this observation. In 2013, the molecular profiling of EC performed by The Cancer Genome Atlas (TCGA) consortium rendered a new molecular classification in four groups: Ultramutated, characterized by pathogenic somatic mutations in the exonuclease domain of the replicative DNA polymerase epsilon (*POLE*); hypermutated/microsatellite unstable group, characterized by microsatellite instability as a result of defective mismatch repair (MSI); Copy Number Low (CNL), composed of *P53* wild-type and *POLE* wild-type, MMR-proficient tumors with relatively low somatic copy number alterations and; Copy Number High (CNH)-Serous-like, characterized by mutations in p53 and extensive somatic copy number alterations[6]. Molecular classification provided a relevant advance in the diagnostic, prognostic, and therapeutic management of EC. Among all molecular alterations identified in EC, those affecting Pten and p53 tumor suppressor genes are among the most frequent ones. Histologically, Pten alterations account for the majority of endometroid carcinomas while most p53 mutated endometrial cancers are serous type, carcinosarcomas and high grade endometrioid carcinomas, but also other histologic types, including low-grade tumors[3]. EC harboring p53 mutations has reliably been associated with the poorest clinical outcomes, with high recurrence risk and particularly high mortality (50-70 % of EC deaths)[1] [2].

Genetically Engineered Mouse Models (GEMMs) that recapitulate the major drivers of EC have been used to model the pathogenesis of this type of neoplasm [7,8]. In the late 1990s, Pten was identified as a frequently mutated gene in ECs[9–11]. In the last decade, genomic landscape of EC confirmed Pten as the most frequently mutated tumor suppressor in EC [3,6]. The role of Pten mutations in the EC development has been confirmed by a genetically engineered mouse models carrying heterozygous deletion of Pten [12–14] or several mouse models for conditional deletion of Pten in the uterus or endometrial epithelium [7]. However, the results provided by mouse models lacking p53 alone or p53 and Pten EC development is more controversial. To date, there are only two mouse models describing the effects of p53 deficiency[15] or combined Pten and/or p53[16] deficiency in EC development. In the first one, mice with endometrium-specific deletion of p53 developed carcinomas representing all type II histological subtypes, including serous, clear cell, undifferentiated EEC and UCS[15]. In contrast in the second one, conditional deletion of endometrial p53 had no phenotype, and double p53 and Pten deletion mice caused invasive EECs, with no signs of UCS, clear cell or serous carcinomas. Therefore, the functional and morphological consequences of co-existing Pten and p53 deficiency in EC are still controversial. Such discrepancies can be explained by differential cellular or cellular Cre activities of the GEMMs used in each study[7]. There is not a Cre expressing mice model for local, tissue specific and temporally controllable expression of Cre recombinase for conditional deletion of the genes of interests restricted to epithelial cell compartment of the uterus without affecting other organs.

Besides the use of Cre based systems to model EC, we have recently demonstrated specific mutation of tumor suppressor genes in the epithelial cells of the endometrium can be achieved by in vivo electroporation of CRISPR/Cas9 components into mouse uterine cavity[17].

Here, we have used three novel methodologies to uncover the role of these two tumor suppressor genes in EC. Using a novel method for intravaginal delivery of TAT-fused CRE recombinase, endometrial xenotransplants, and CRISPR/cas9 technology we have analyzed the phenotype of simultaneous epithelial cell-specific deletion of Pten and p53. Our results demonstrate that epithelial specific simultaneous deletion of Pten and p53 in epithelial cells, without affecting stromal or myometrial cells, is sufficient to cause EMT that triggers the development of metastatic high-grade EEC and UCS, resembling those observed in human ECs.

## MATERIAL AND METHODS

**Mouse models.** Mice were housed in a barrier facility and pathogen-free procedures were used in all mouse rooms. Animals were kept in a 12-hour light/dark cycle at 22°C with *ad libitum* access to food and water. All procedures performed in this study followed the National Institute of Health Guide for the Care and Use of Laboratory Animals and were compliant with the guidelines of Universitat de Lleida. Conditional Pten knock-out mice (Pten floxed) (C;129S4-*Ptentm1Hwu*/J, hereafter called PTEN^fl/fl^); conditional p53 knock-out mice (p53 floxed) (FVB.129P2-*Trp53^tm1Brn^*/Nci), Cre:ER^T^ mice (B6.Cg-Tg(CAG-CRE/Esr1* 5Amc/J) and the reporter mT/mG mice (B6.129(Cg)-*Gt(ROSA)26Sortm4(ACTB-tdTomato,-EGFP)Luo*/J) were obtained from the Jackson Laboratory (Bar Harbor, ME, USA). Genotyping primers and PCR conditions are specified in Supplementary methods (Table SM1). Immunodeficient female SCID hr/hr mice were bred in Universitat de Lleida animal housing facility.

### Cloning, expression and isolation of recombinant TAT-CRE

Cre recombinase DNA was amplified by PCR using pCAG-Cre (Addgene #13775) as a template. A 6X histidine tag (His), nuclear localization sequence and TAT sequence were added to the 5’ end of Cre DNA by PCR. The resulting His-TAT-NLS-Cre DNA fragment was cloned in pLATE11 vector using the aLICator Ligation Independent Cloning and expression Kit (Thermo) following manufacturer’s instructions. The resulting plasmid was transformed in Escherichia Coli BL21 (DE3) comnpetent cells (MERCK). Expression of TAT-Cre was induced growing transformed cells in LB Overnight Express™ Instant medium (Novagen) overnight at 30°C. Bacteria were collected by centrifugation at 8000 r.pm. and lysed in of 50 mM Tris base (FISHER) at pH7.5, 1 M NaCl (Merck), 10% glycerol, 0.1% Triton-X (Sigma-Aldrich) and 20 mm Imidazole (SIGMA), and cOmplete™ Protease Inhibitor Cocktail (Roche), 5mM β-Mercaptoethanol (Bio-Rad), 1 mg/mL lysozyme (Roche). Lysate was allowed to settle for 30 min on ice and sonicated for 1 minute (20 second intervals on ice). To purify His-tagged TAT-Cre lysates were incubated with Cobalt HisPur™ resin (Thermo) for 1 hours at 4°C and washed three times with lysis buffer. TAT-Cre was eluted by incubating resin was incubated with lysis buffer supplemented with 300 mM imidazole. Eluted Tat-Cre was concentrated by centrifugation at 4000 rpm for 15 minutes at 4°C using and Amicon® Ultra-15 Centrifugal Filter Unit. Concrentrated TAT-Cre was diluted 1:1 in PBS supplemented with 20mM HEPES, pH 7.4, 50% glycerol and 500mM NaCl and stored at −20°C. The resulting volume containing TAT-Cre was quantified using NanoDrop™ spectrophotometer obtaining a concentration of 17.6 mg/mL. Purity and size of TAT-Cre was also checked by loading a polyacrylamide gel with the equivalent to 3.2 µg of purified TAT-Cre Supplementary Methods (Figure SM1).

### Isolation and culture of mouse skin fibroblast

Tail and ear fragments from mT/mG mice were dissected and incubated in 2-3 hours at 37 °C in 950 µL of Dulbecco’s modified Eagle’s medium (DMEM, Invitrogen), supplemented with 10% inactivated fetal bovine serum (FBSi, Invitrogen), 1 mmol/L HEPES (Sigma -Aldrich, San Luis, MO), 1 mmol/L sodium pyruvate (Sigma-Aldrich), 1 % penicillin/streptomycin (Sigma-Aldrich) and 0.1 % Amphotericin B (Invitrogen) (supplemented DMEM) and 50 µL Collagenase Type IA 200 mg/mL (Worthington Biochemical Corporation Lakewood, NJ). Digested skin fragments were filtered through sterile Cell Strainer (70 µM Nylon Mesh, Fisherbrand, Hampton, NH). Cell strainer was washed with supplemented DMEM medium to recover most of digested cells. Isolated fibroblasts were centrifuged, resuspended in complete medium and plated in P100 cultures plates. Twenty-four hours after plating, medium was changed for fresh supplemented DMEM medium.

### Isolation of epithelial and mesenchymal cells from the mouse uterus and establishment of three-dimensional organoid cultures of endometrial epithelial cells

Isolation of mouse endometrial epithelial cells was performed as previously described [18]. Mice were sacrificed by cervical dislocation, and uterine horns were dissected from electroporated mT/mG mice. Uteri were washed with Hanks Balanced Salt Solution (HBSS) (Invitrogen) and chopped in 3- to 4-mm-length fragments. Uterine fragments were digested with 1% trypsin (Invitrogen) in HBSS for 1 hour at 4°C and 45 minutes at room temperature. Trypsin digestion was stopped by addition of DMEM containing 10% fetal bovine serum (Invitrogen). After trypsin digestion, epithelial sheets were squeezed out of the uterine pieces by applying gentle pressure with the edge of a razor blade. Epithelial sheets were washed twice with PBS and resuspended in 1 ml of DMEM/F12 (Invitrogen) supplemented with 1 mmol/L HEPES (Sigma-Aldrich), 1% penicillin/streptomycin (Sigma-Aldrich), and Fungizone (Invitrogen) (basal medium). Epithelial sheets were mechanically disrupted in basal medium by pipetting 50 times through a 1-ml tip, until clumps of cells are observed under the microscope. Cells were diluted in basal medium containing 2% dextran-coated charcoal-stripped serum (DCC) (HyClone Laboratories, Logan, UT) and plated into culture dishes (BD Falcon, Bedford, MA). Cells were cultured for 24 hours in an incubator at 37°C with saturating humidity and 5% CO_2_. Twenty-four hours after platting, cells were washed with HBSS and incubated with trypsin/EDTA solution (Sigma) for 5 min at 37°C. Trypsin was stopped by adding DMEM 10 % FBS and clumps of 2-8 cells were obtained. Cells were centrifuged at 1000 rpm for 3 minutes and diluted in basal medium containing 3 % of Matrigel to obtain 4 x 10^4^ cell clumps/ml. For immunofluorescence, cells were seeded in a volume of 40 μl/well in 96 well plates (black with micro-clear bottom) (Greiner Bio-one). For western blotting, cells were placed in a volume of 200 μl in 24-well plates (BD Biosciences). In all cases, 24 hours after plating medium was replaced by Basal medium supplemented with 5 ng/ml EGF and 1/100 dilution of Insulin-Transferrin-Sodium Selenite (ITS) Supplement (Invitrogen) and 3 % of fresh Matrigel. Medium was replaced every 2-3 days.

### Tamoxifen treatments

Tamoxifen (Sigma-Aldrich T5648, St Louis, MO) was dissolved in 100% ethanol at 100 mg/ml. Tamoxifen solution was emulsified in corn oil (Sigma-Aldrich C8267) at 10 mg/ml by vortexing. Adult mice (4–5 weeks old) were given a single intraperitoneal injection of 0.5 mg of tamoxifen emulsion (30–35 μg per mg body weight).

### Glandular perimeter measurements

Images of endometrial epithelial spheroids were captured and digitized with a confocal microscope (Fluoview FV1000, Olympus). Epithelial perimeter analysis was processed by image analysis software (ImageJ version 1.46r; NIH, Bethesda, MD, USA), generating binary images of the spheroids as previously described[19].

### BrdU labeling

Organoids cultures were incubated with 3 ng/ml of 5-bromodeoxyuridine (BrdU, Sigma-Aldrich) during 15 h and then fixed with 4 % paraformaldehyde for 20 min. DNA denaturation was performed with 2 mol/l HCl for 30 min. Afterwards, neutralization was done with 0.1 mol/l Na_2_B_4_O_7_ (pH 8.5) for 2 min and after rinsed three times with PBS. Subsequently, block cells in PBS solution containing 5% horse serum, 5% FBS, 0.2% glycine and 0.1% Triton X-100 for 1 h. Next, cells were incubated in anti-BrdU primary monoclonal antibody (DAKO, Glostrup, Denmark), and fluorescein isothiocyanate-conjugated anti-mouse secondary antibody. Nuclei were counterstained with 5 mg/ml Hoechst 33258 and cells were visualized under confocal microscope. Cells were visualized under a confocal microscope. BrdU-positive nuclei were scored and divided by the total number of cells (visualized by Hoechst staining). The results are expressed as a percentage of BrdU-positive cells.

### Measurement of the nuclear area

Measurement of the nuclear area was performed with the QuPath 0.1.2 software. On histological sections stained with hematoxylin-eosin, of subcutaneous tumors of epithelial endometrial cells. The region of interest (ROI) was defined as tumor epithelial cells, excluding extracellular matrix regions and mesenchymal cells. Through an algorithm of the QuPath software, the cells that fell within the standard marked as epithelial cell were detected and a colorimetric scale from purple to yellow was applied that corresponded to the smallest nuclear area (10 µm2) to the largest (210 µm2), respectively.

### In vitro TAT-Cre assay

In vitro TAT-Cre recombinase was performed by diluting 0,05 µg or 0.1µg or 0.2µg of homemade TAT-Cre in a final volume of 10µL of Opti-MEM reduced serum medium (GIBCO). Next, 2 µg of pCA-mTmG plasmid (#26123, Addgene) was added to diluted TAT-Cre and incubated at 37°C for 1 hour. Digestions were resolved on a 1% agarose gel.

### mRNA extraction, sequencing, and Gene Set Enrichment Analysis (GSEA)

Total RNA from organoid cultures was isolated with the SurePrep™ TrueTotal™ RNA Purification kit (Fisher), following the manufacturer’s instructions. The RNA extracts were quantified with the NanoDrop® (ND-1000 UV/Vis Spectophotometer, Nanodrop Technologies) and stored frozen at −80°C.

Total RNA was submitted to whole transcriptome analysis (RNA-seq) at National the Centre of Genomics (CNAG)-Center for genomic Regulation (CRG-Barcelona). Data from RNA-seq was subjected to Gene Set Enrichment Analysis (GSEA) The gene set shown in this work has the pertinent statistical values to be considered as significantly associated with one of the conditions (p<0.05; FDR<0.25).

### In vitro and in vivo delivery of TAT-Cre

For in vitro delivery of TAT-Cre to mouse fibroblasts, endometrial epithelial cell cultures in two- or three-dimensions cell culture medium was replaced by Opti-MEM™ reduced serum medium (GIBCO) containing recombinant TAT-Cre indicated in each experiment. After 16 hours of incubation, the TAT-Cre containing medium was replaced by fresh medium corresponding to each cell type.

For in vivo intrauterine delivery of TAT-Cre, mice were anesthetized with 2 % isoflurane by inhalation. A longitudinal incision in the skin and peritoneal wall in the abdomen was performed using sterile dissection scissors and forceps. Uterus was located and pulled out of the abdominal cavity. Once uterus was exposed, 10 µL of recombinant Tat-Cre at 8.8 µg/µL supplemented with Fast Green FCF (Sigma) to facilitate visualization of TAT-Cre delivery were intravaginally injected to each uterine horn using a micropipette tip (Figure 4A). Uterus was carefully reintroduced back into the peritoneal cavity, and the skin and peritoneal wall was closed with a wound stapler (AutoClip System 12020-09, Fine Scientific Tools, FST, Poznań, Poland).

### Tissue processing and immunohistochemistry analysis on paraffin sections

Animals were euthanized and uteri were collected, formalin-fixed O/N at 4 °C and embedded in paraffin for histologic examination. Three µm sections of the paraffin blocks were dried for one hour at 80 °C, dewaxed in xylene, rehydrated through a graded ethanol series, and washed with phosphate-buffered saline (PBS). Antigen retrieval was performed in EnVision™ FLEX, High pH or Low pH solution (DAKO, Glostrup, Denmark) for 20 minutes at 95°C, depending on the primary antibody. Endogenous peroxidase was blocked by incubating slides with 3% H2O2 solution. After three washes with phosphate-buffered saline (PBS) (DAKO), primary antibody was incubated for 30 minutes at room temperature, washed in PBS and incubated with secondary antibody. A peroxidase (HRP)-conjugated secondary antibody or a biotin conjugated secondary antibody plus peroxidase-linked streptavidin were used depending on the primary antibody used. The reaction was visualized with the EnVision Detection Kit (DAKO) using diaminobenzidine chromogen as substrate. Sections were counterstained with Harris haematoxylin. Antibodies used in this study and immunohistochemistry conditions are summarized in Supplementary Methods (Table SM2)

### Immunofluorescence

Immunofluorescence was performed as previously described [18,19]. Cultures were fixed with 4% paraformaldehyde in PBS for 15 minutes at room temperature (RT), washed twice with PBS, permeabilized with 0,2 % triton X-100 in PBS for 10 minutes and blocked with PBS containing 2% Horse serum, 2% Bovine Serum Albumin (BSA) 0,2% Triton X-100 and incubated overnight at 4 °C with the indicated dilutions of antibodies listed in Supplementary Methods (Table SM3) After one day, cells were washed twice with PBS and incubated with PBS containing a 5 μg/ml of Hoechst dye and 1/500 dilution of Alexa Fluor secondary anti-mouse or anti-rabbit antibodies for 2 hours at RT in the case of 2D cultures or overnight at 4 °C for 3D cultures. For double immunofluorescence staining, cells were incubated with the second round of primary and secondary antibodies. We would like to point out that in all double immunofluorescence stains, first and second primary antibodies were from different isotope. Immunofluorescence staining was visualized and analyzed using a confocal microscopy Olympus FluoView™1000 (Olympus).

### Western blot

Western blot was performed as previously described[19,20]. Briefly, 2D endometrial monolayers or endometrial organoid 3D cultures stimulated for the indicated periods of time, were washed with HBSS and incubated with trypsin/EDTA solution for 5 min at 37 °C. Incubation with trypsin was done to allow us to separate the glandular structures from Matrigel. Trypsin was stopped by adding DMEM 10 % FBS and the cells were lysed with lysis buffer (2 % SDS, 125 mM Tris-HCL pH6.8). Relative protein concentrations were determined loading an 8% acrylamide gel, transferred to PVDF membranes and blotted with anti-tubulin antibody. Equal amounts of proteins were subjected to SDS-PAGE and transferred to PVDF membranes (Millipore, Bedford, MA). Non-specific binding was blocked by incubation with TBST (20 mM Tris-Hcl pH7.4, 150 mM NaCl, 0.1 % Tween-20) plus 5 % of non-fat milk. Membranes were incubated overnight at 4 °C with primary antibodies listed in Supplementary Methods (Table SM4). The procedure was followed by 1-hour incubation with secondary antibody 1/10000 in TBST at RT. Signal was detected with Immobilon Forte Western HRP Substrate (EMD Millipore Corporation, Burlington).

### Subcutaneous xenotransplants

Mice with the indicated genotypes (Figure 7) were intraperitoneally injected with tamoxifen to induce Pten and/or p53 ablation in endometrial epithelial cells. The day after injection, endometrial epithelial cells from tamoxifen injected mice and uterine mesenchymal cells from C57BL/6 mice were isolated and cultured as described in “Isolation of epithelial and mesenchymal cells from the mouse uterus and establishment of three-dimensional cultures of endometrial epithelial cells” section. Twenty-four hours after plating, wild type mesenchymal cells and epithelial cells of the indicated genotypes were incubated with trypsin/EDTA solution (Sigma) for 5 min at 37°C, resuspended in DMEM 10% FBS to stop trypsin activity, washed with PBS and resuspended in DMEM/F12 medium supplemented with 1 mM HEPES, 1% penicillin/streptomycin, 0.1% amphotericin B, 2% DCC, and 20% Matrigel. Resuspended epithelial and stromal cells were mixed 1:1 and 100 µL of cells were subcutaneously injected each flank of SCID females between 8-12 weeks of age.

Tumor growth was monitored every 2 weeks with a digital caliper and tumor volume was calculated according to the formula: TV (tumor volume) = (Dxd2)/2, where D corresponds to the large diameter of the tumor and d, to the small one. Mice were sacrificed by cervical dislocation when tumors reached 1 cm^3^. Tumors are collected for macroscopic and immunohistology analysis.

### In vivo CRISPR/Cas9 RNP electroporation into mouse uterine cavity

In vivo Cas9 RNP electroporation was performed as recently described[17]. Briefly, recombinant Streptococcus pyogenes Cas9. (Alt-R® S.p Cas9 Nuclease V3, IDT) was mixed at equimolar concentration with crRNA:tracrRNAs (IDT) and diluted in Opti-MEM (Invitrogen, Waltham, MA) at a final concentration of 6 µM in 5 µL (0.5 µL of 61 µM Cas9, 1.5 µL of 20 µM crRNA:tracrRNA and 3 µL of Opti-MEM). Mixture was incubated for 20-30 minutes at room temperature to allow Cas9-crRNA:tracrRNA RNP formation (Cas9-RNP).cRNA sequences (including underlined PAM) targeting Pten and p53 were 5’-AAUUCACUGUAAAGCUGGAA-3’ and 5’-GGAGUCUUCCAGUGUGAUGA-3’, respectively (IDT).

Mice were anesthetized with 2 % isoflurane by inhalation. A longitudinal incision in the skin and peritoneal wall in the abdomen was performed using sterile dissection scissors and forceps. Uterus was located and pulled out of the abdominal cavity. Once uterus was exposed, 5 µL of 6 µM RNP supplemented with Fast Green FCF (Sigma) to facilitate visualization of RNP complexes delivery were injected to each uterine horn using a Hamilton Neuros syringe (Hamilton, Reno, NV). For the injection of more than one Cas9-RNP complexes, 5 µl of mixture in 1:1 ratio of each Cas9-RNP was injected per uterine horn. Using a BTX830 square electroporator (BTX, Hawthorne, NY), 4 pulses of 50 mV for 50 msec spaced by 950 msec were applied to the injected uterine horn. This protocol was repeated opposing the orientation of tweezers and performed along all the entire uterine horn. The tweezers used were Platinum Tweezertrode, 5 mm Diameter (BTX). Once electroporated, uterus was carefully reintroduced back into the peritoneal cavity, and the skin and peritoneal wall was closed with a wound stapler (AutoClip System 12020-09, Fine Scientific Tools, FST, Poznań, Poland).

### Amplicon-Next Generation Sequencing and data analysis

Electroporated uteri were dissected and opened longitudinally to expose the endometrial epithelial cells of the uterine lumen. Under a fluorescence stereoscopic microscope (Nikon Eclipse Ts2R), luminal epithelial areas displaying green cells were rinsed with HBSS (Invitrogen) and scraped with a scalpel blade. Scraped sheets of epithelial cells were collected rinsed with PBS based on a series of centrifugations. Cell pellet was processed to extract the genomic DNA from the endometrial epithelial cells using the NZY Tissue gDNA Isolation Kit (NYZTech), following the manufacturers’ instructions. Genomic DNA was amplified by PCR using primers flanking RNP-targeted region. Primer sequences and PCR conditions for amplification of DNA fragments are specified in Supplementary Methods (Table SM5). PCR amplicons were resolved in agarose gels, purified using the NZYGelpure kit (NZYTech) following manufacturers’ instructions and quantified with a NanoDrop spectrophotometer. Next Generation Sequencing (NGS)-Amplicon sequencing was carried out by Genewiz (Azenta Life Sicences, Chelmsford, MA) company. Raw FASQ archives containing NGS-amplicon sequences were submitted to bioinformatic analysis to detect the presence of indels using Crispresso2 [21] and Cas-Analyzer [22] software.

## RESULTS

### In vivo Tamoxifen-inducible deletion of Pten and P53 leads to the development of hyperplasia and non-invasive intraepithelial neoplasia

First, to address the effects of double Pten and p53 deficiency in EC histopathology, we aimed to generate an in vivo tamoxifen-inducible mouse model to study the effects of double Pten and p53 deletion in the development and progression of EC: For this purpose, we crossed our previously reported mouse model for tamoxifen-inducible Pten deletion (Cre:ER^T^; Pten^f/f^)[23] with conditional p53 knockout mice (p53^f/f^). After appropriate breeding protocol, we obtained mice carrying both Pten and p53 floxed alleles and the tamoxifen inducible Cre recombinase (Cre:ER^T+/-^; Pten^f/f^; P53^f/f^),and all the control combinations of wild type and floxed Pten and p53 alleles (Supplementary Figure 1A). Once obtained the desired genotypes, five-week-old females were intraperitoneally injected with a single dose of tamoxifen (0,5mg/Kg) and sacrificed 6 weeks later. To examine the histopathological effects of double Pten and p53 deletion over time, five-week-old females were injected with tamoxifen and mice were sacrificed in three-time brackets 1-2 weeks, 3-4 weeks or 5-6 weeks post-injection. No endometrial lesions were observed in mice sacrificed after 1-2 weeks pos-tamoxifen injection irrespective of the genotype (Supplementary Figure 1B). In the second time period, 30% of the Cre:ER^+/-^; Pten^f/f^; p53^+/+^, Cre:ER^+/-^; Pten^f/f^; p53^f/+^ and Cre:ER^+/-^; Pten^f/f^; p53^f/f^ mice developed complex hyperplasia, while 50-55% of the mice displayed hyperplasia with complex atypia and endometrioid adenocarcinomas limited to the endometrium. In the third group, virtually all Cre:ER^+/-^; Pten^f/f^ mice presented low-grade EC regardless p53 genotype (Supplementary Figure 1B). Thus, no statistically significant histopathological differences were observed among the experimental genotypes presenting Pten ablation. In addition, we observed no differences in the expression of the proliferation marker Ki-67 among endometrial lesions of Cre:ER^+/-^; Pten^f/f^; p53^+/+^, Cre:ER^+/-^; Pten^f/f^; p53^f/+^ and Cre:ER^+/-^; Pten^f/f^; p53^f/f^ (Supplementary Figure 1C). These results indicate that at the periods of time analyzed, p53 had no effect on endometrial lesions caused by Pten loss.

To further investigate the effect of p53 deficiency on endometrial lesions caused by Pten loss, we extended period of study after tamoxifen administration. However, all Pten or double Pten/p53 deficient mice presented severe respiratory difficulties, lethargy and anorexia that appeared earlier in Pten/p53 double knock-out (6 weeks) than in Pten knock-out (8 weeks). Kaplann-meyer plot demonstrated decreased survival of Pten knock-out mice lacking either one or two p53 alleles (Supplementary Figure 1D). The cause of premature death of Pten-deficient mice was previously described[23]. In case of double Pten/p53 double knock-out, we performed a gross histopathological study that revealed proliferative lesions in multiple organs such as colon adenomas, hepatocellular dysplasia, severe thyroid hyperplasia, or lymphoma (Supplementary Figure 2), thus limiting the study of double Pten and p53 deletion in EC.

### Double Pten and p53 deletion triggers EMT and invasive phenotype of three-dimensional endometrial organoids

The genomic profiling demonstrated that EC harboring Pten and p53 mutations are associated with a highly aggressive behavior[6], but the above presented model did not recapitulate such feature. Our previous studies demonstrate that three dimensional (3D) cultures of endometrial organoids recapitulate endometrial alterations caused by Pten deficiency[18]. Tamoxifen-induced Pten ablation in endometrial organoids isolated from Cre:ERT^+/-^; Pten^f/f^ mice resulted in increased proliferation and resistance to apoptosis, but no signs of invasive phenotype were observed [19]. Therefore, to insight into effects of p53 and concurrent Pten/p53 deletion in endometrial organoids, endometrial epithelial cells were isolated from Pten^+/+^; p53^+/+^(WT)^;^ Pten^f/f^; p53^+/+^(PTENKO), Pten^+/+^; p53^f/f^ (p53KO), Pten^f/f^; p53^f/f^ (dKO) mice and were treated with tamoxifen to induce Pten and/or p53 ablation and we examined the effects on proliferation, invasiveness and EMT. Double Pten/p53 deficiency caused a further on increase glandular perimeter (Figure 1A) and BrdU incorporation (Figure 1B) of 3D organoids compared to those lacking Pten, suggesting that p53 deficiency enhances proliferation of Pten deficient cells. To analyze invasiveness, we evaluated the growth of invadopodia into the matrigel surrounding endometrial organoids. Double Pten/p53 deficiency caused a marked increase in the number of organoids displaying invasive invadopodia over organoids lacking only Pten (Figure 1C). We further analyzed whether such invasive phenotype was associated with EMT by immunofluorescence analysis of Cytokeratin 8 and vimentin (Figure 1D). Cells displaying invasive phenotype exhibited loss of the epithelial marker cytokeratin and expression of the mesenchymal marker vimentin, suggesting that these cells were undergoing an EMT process. Finally, to demonstrate a EMT signature of double Pten/p53 organoids, we performed a mRNA-seq analysis of Pten and double Pten/p53-deficient organoids. Bioinformatic study of mRNA-seq data using GSEA revealed an increase of the hallmark EMT on double Pten/p53·organoids over organoids with Pten deletion (Figure 1E).

**Figure 1.**
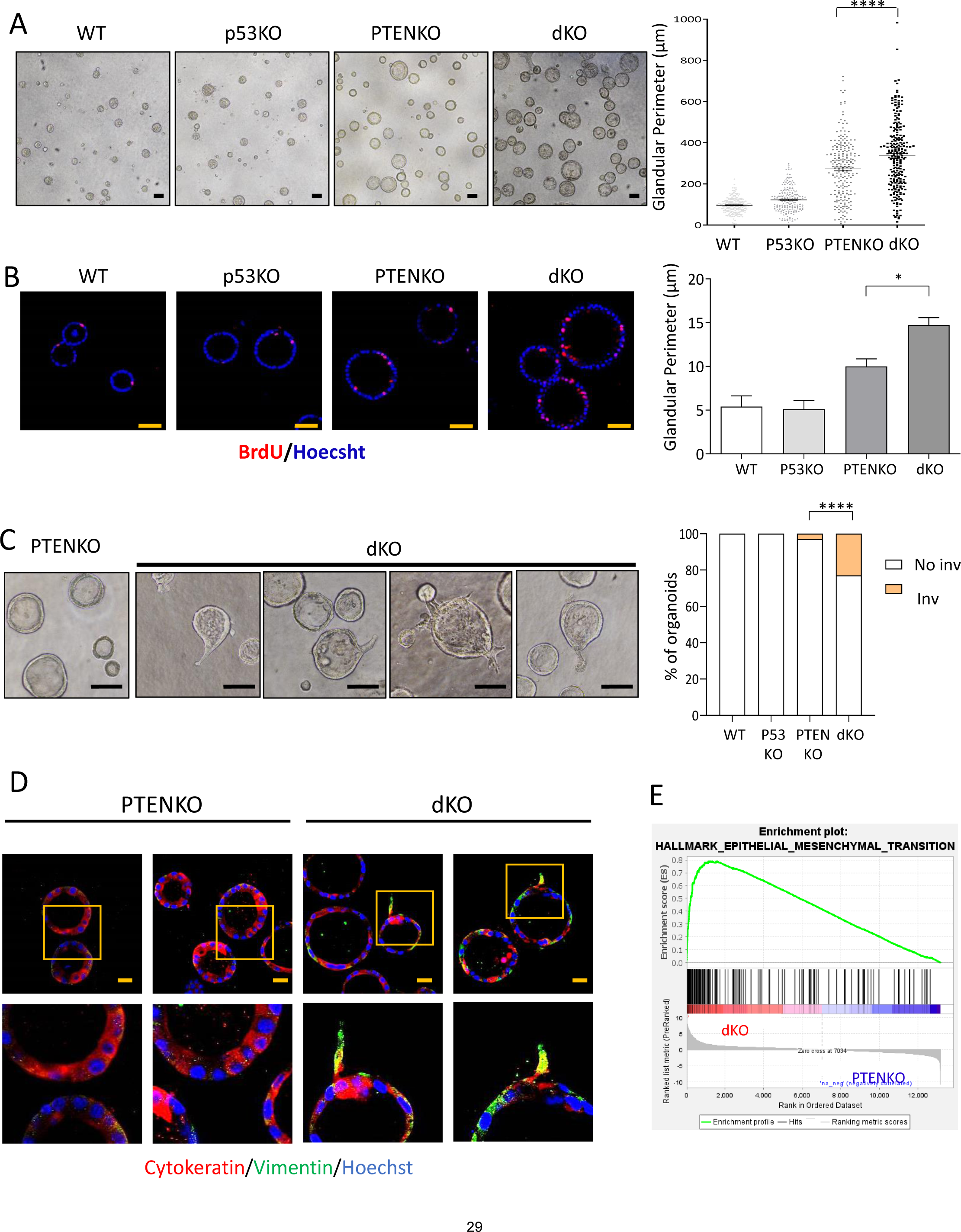
Loss of Pten and p53 increases proliferation, invasive phenotype and EMT in endometrial organoids. **(A)** Representative phase contrast images and quantification organoid cultures obtained from Pten^+/+^; p53^+/+^ (WT), Pten^+/+^; p53^f/f^ (p53KO), Pten^f/f^; p53^+/+^ (PTENKO), and Pten^f/f^; p53^f/f^ (dKO) mice. Scale bars: 100µm. n=3 independent experiments. Error bars represent mean ± s.e.m. ****p<0.0001, according to the Bonferroni multiple comparison test **(B)** Representative images and quantification of BrdU immunofluorescence of organoid cultures of epithelial cells isolated from from Pten^+/+^; p53^+/+^ (WT), Pten^+/+^; p53^f/f^ (p53KO), Pten^f/f^; p53^+/+^ (PTENKO), and Pten^f/f^; p53^f/f^ (dKO) mice. Scale bars: 50µm. n=3 independent experiments. Values are represented as the mean ± s.e.m. p*<0.05, one-way ANOVA analysis, followed by a Tukey multiple comparison test. **(C)** Left, representative phase contrast images of one Pten deficient (PTENKO) or three double PTEN/p53 deficient (dKO) endometrial organoids showing invadopodia. Right, quantification of the number of glands presenting invadopodia in organoids from Pten^+/+^; p53^+/+^ (WT), Pten^+/+^; p53^f/f^ (p53KO), Pten^f/f^; p53^+/+^ (PTENKO), and Pten^f/f^; p53^f/f^ (dKO) mice. n=3 independent experiments. Hundred organoids per genotype were scored. ****p<0.0001, according to Fisher’s exact test with corrected standards. Scale bars: 100µm. **(D)** Representative vimentin and cytokeratin immunofluorescence on organoid cultures obtained from Pten^f/f^ or p53^+/+^ (PTENKO), and Pten^f/f^; p53^f/f^ (dKO) mice. Organoids were stained with Hoechst to visualize nuclei. **(E)** Gene Set Enrichment analysis (GSEA) transcriptomic analysis of RNA sequencing results obtained from RNAs isolated from Pten^f/f^; p53^+/+^ (PTENKO) or Pten^f/f^; p53^f/f^ (dKO) organoids.

### TAT-mediated delivery of Cre recombinase induce recombination of LoxP-flanked alleles in vitro and in vivo

Data obtained from the study in endometrial organoids, suggested that double Pten/p53 deficiency could lead EMT and development of invasive endometrial carcinoma. However, the abovementioned survival problems associated with the development of extrauterine pathologies in tamoxifen induced Pten/p53 double KO, limited in vivo study its role in endometrial carcinogenesis. To override such limitations, we aimed to develop a strategy to induce localized and inducible conditional ablation of floxed alleles in the uterine cavity without affecting other organs. To date, there is no Cre-expressing mouse models allowing both conditional and inducible deletion of endometrial epithelial cells without affecting other tissues[7]. Therefore, we proceed to local administration of recombinant Cre recombinase into the mouse uterus. To achieve intracellular delivery of Cre into the uterine cells, we constructed and produced a TAT-fused recombinant Cre recombinase. Recombinant TAT-Cre was produced, isolated, and purified in E. Coli as described in the methods section. Once obtained, preservation of TAT-Cre nuclease activity, was analyzed by an in vitro assay. Increasing doses of TAT-Cre were incubated with the pCA-mT/mG reported plasmid and the reaction was resolved in agarose gel (see material and methods for details). TAT-Cre caused the cleavage of a DNA fragment corresponding to the size LoxP-flanked mT sequence (Supplementary Figure 3A-3B). Having demonstrated that TAT fusion did not disrupt Cre nuclease activity, we analyzed its ability to translocate and cause genomic recombination of LoxP sites on living cultured cells, tested whether it was able to translocate across cell membranes and cause efficient recombination of floxed alleles. For this purpose, we took advantage of the mT/mG reporter mice. These mice possess *LoxP* -flanked membrane-targeted tdTomato (mT) cassette and express strong red fluorescence in all tissues and cell types [24]. Breeding mT/mG reporter mice to Cre recombinase expressing mice, results in mT cassette deletion in the *Cre* expressing tissues, allowing expression of the downstream membrane-targeted EGFP (mG) cassette. We first determined the activity and optimal concentration of TAT-Cre in vitro. Monolayer (Figure 2A) or organoid cultures (Figure 2B) of mouse endometrial epithelial cells and skin fibroblasts (Supplementary Figure 3C) isolated from mT/mG mice were incubated with increasing concentrations of TAT-Cre and the number of red (expressing mT, non-recombined) and green cells (expressing mG, recombined) was scored as readout of recombination activity. In all three cultures, TAT-Cre caused a dose-dependent increase of the number of mG expression cells, reaching the maximum values at 176-352µg/mL concentration.

**Figure 2.**
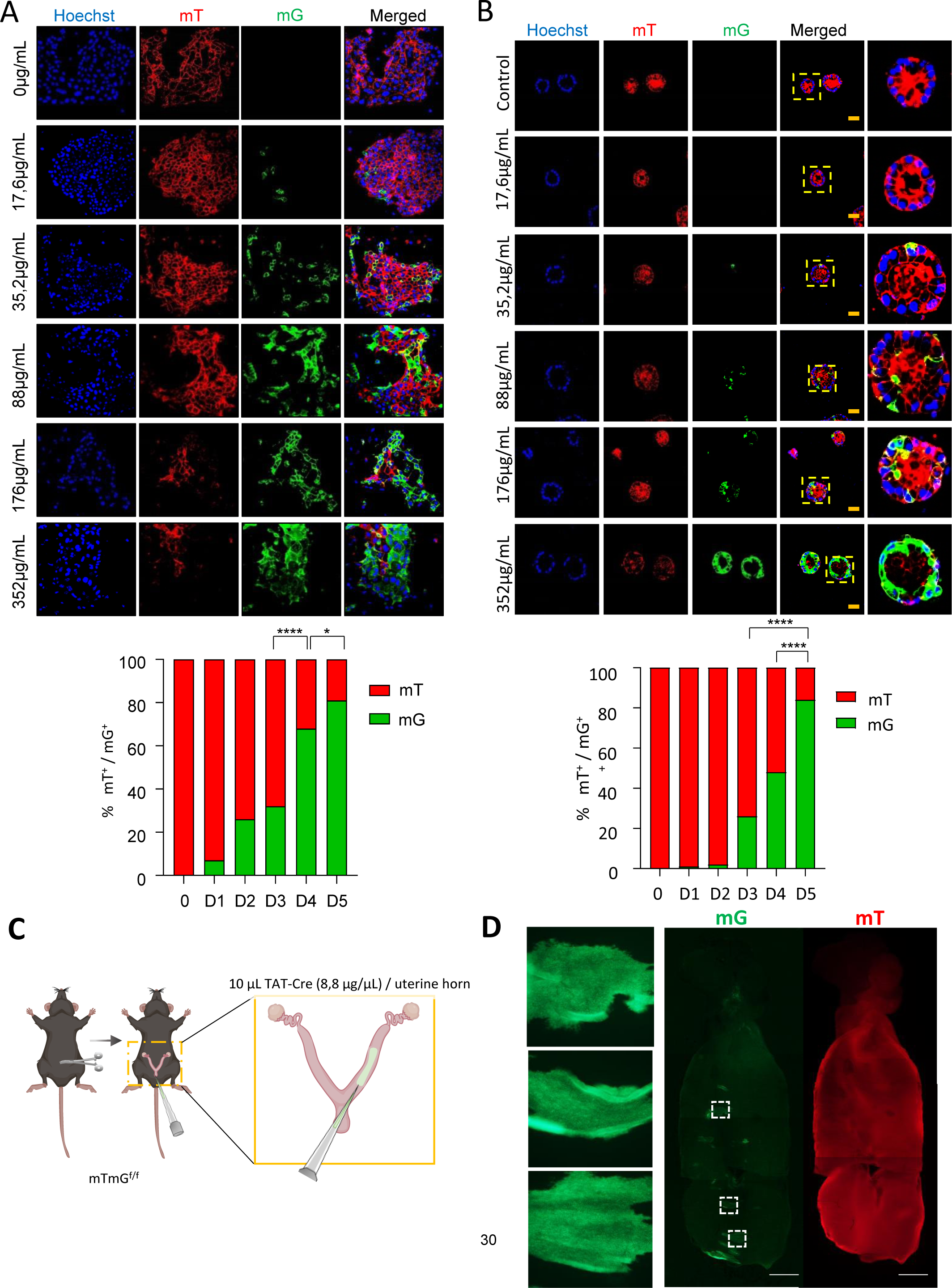
Recombinant TAT-Cre exhibits recombinase activity on endometrial epithelial cells in vitro and in vivo. **(A)** Representative images and quantification of tdTomato positive (mT) or GFP positive (mG) monolayer endometrial epithelial cells isolated from mT/mG^f/f^ mouse ears 4 days after being treated with the following TAT-CRE concentrations: 0µg/mL (0), 17.6µg/mL (D1), 35.2µg/mL (D2), 88µg/mL (D3), 176µg/mL (D4) and 352µg/mL (D5). Data come from n=3 independent experiments. *p<0.5 ****p<0.0001, using a one-way ANOVA analysis, followed by a Bonferroni multiple comparison test. **(B)** Representative images and quantification of tdTomato positive (mT) or GFP positive (mG) organoids of endometrial epithelial cells isolated from mT/mG^f/f^ mouse ears 4 days after being treated with the following TAT-CRE concentrations: 0µg/mL (0), 17.6µg/mL (D1), 35.2µg/mL (D2), 88µg/mL (D3), 176µg/mL (D4) and 352µg/mL (D5). Data come from n=3 independent experiments. *p<0.5 ****p<0.0001, using a one-way ANOVA analysis, followed by a Bonferroni multiple comparison test. **(C)** Diagram depicting intrauterine administration of 88 µg of recombinant TAT-CRE per uterine horn of mT/mG^f/f^ mouse. **(D)** Right, representative images images obtained from red (tdTomato) and green (mGFP) fluorescence at 4X of mT/mG^f/f^ endometria 4 months after intravaginal administration of 88µg of recombinant TAT-CRE. Scale bar at 2mm. Left, 20x magnification of the indicated framed areas where the membranous mGFP labeling is appreciated in endometrial cells 4 month after TAT-Cre-mediated gene ablation.

**Figure 3.**
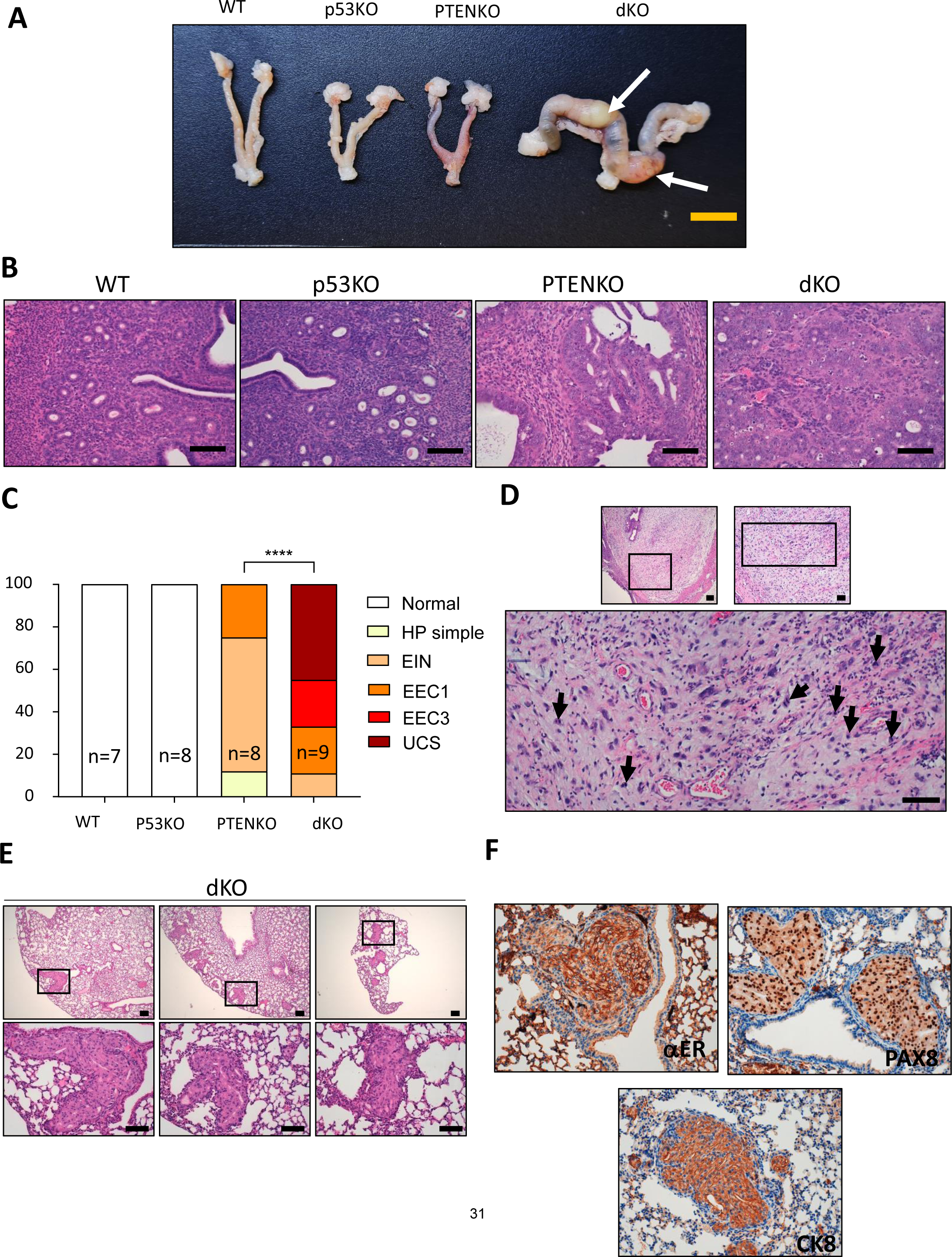
Loss of Pten and p53 induced by intravaginal administration of TAT-CRE results in high-grade endometrial carcinomas and UCS. **(A)** Representative macroscopic images of the uteri of Pten^+/+^ mice; p53^+/+^ (WT), Pten^+/+^; p53^f/f^ (p53KO), Pten^f/f^; p53^+/+^ (PTENKO), and Pten^f/f^; p53^f/f^ (dKO) at 4 months after intravaginal administration of 88µg of TAT-CRE per uterine branch. Representative macroscopic image of a dKO mouse showing enlargement of the uterus and uterine bulges (white arrows). Scale bar: 1cm. **(B)** Representative hematoxylin-eosin staining of endometria from WT, P53KO, PTENKO and dKO mice. Images at 20x. Scale bar: 100µm. **(C)** Quantification of endometrial lesions for the indicated groups of mice. ****p<0.0001, according to the Xi^2^ test, followed by Fisher’s exact test. EIN (Endometrial Intraepithelial Neoplasia), EEC-1 (Grade 1/Low Grade Endometrioid Endometrial Carcinoma), EEC-3 (Grade 3/High Grade Endometrioid Endometrial Carcinoma), UCS (Uterine Carcinosarcoma). **(D)** UCS arised from Pten/p53 deficient endometrial cells display a high mitotic rate and marked nuclear pleomorphism. Representative images of hematoxylin-eosin staining of one of the carcinosarcomas generated at 4 months from the gene deletion induced by intravaginal administration in Pten^f/f^ mice; p53^f/f^ (dKO) of 88 µg of homemade TAT-Cre per uterine branch. Images at 4x, 10x and 20x. Sacale bars: 200µm, 100µm and 100µm, respectively. Black arrows indicate mitoses found in the sarcomatous component of UCS**. (E)** Representative images of hematoxylin-eosin staining of three lung regions showing the micrometastases grown 4 months after the gene deletion induced by intravaginal administration of 88 µg of TAT-CRE per uterine branch in Pten^f/f^ mice; p53^f/f^ (dKO). Images at 4x and 20x. Scale bar: 200µm and 100µm, respectively. **(F)** Representative images of cytokeratin 8 (CK8), estrogen receptor alpha(αER), and Paired Box 8 protein 8 (PAX8) immunostaining of lung micrometastases grown 4 months after the gene deletion induced by intravaginal administration of 88 µg of TAT-Cre per uterine branch in Pten^f/f^ mice; p53^f/f^ (dKO). Images at 20x. Scale bar: 100µm.

Once demonstrate the ability of TAT-Cre to penetrate across membranes and induce LoxP recombination in endometrial epithelial cells in vitro, we tested its activity in vivo. For this purpose, 10 µL of recombinant TAT-Cre at 8.8 µg/µL were intravaginally injected per uterine branch of mTmG mice (Figure 2C). Mice were sacrificed 4 months after injection and the presence of red-to-green fluorescence was analyzed as readout of TAT-Cre activity. Microscopic evaluation of the endometria, revealed the presence of positive areas for green fluorescence, suggesting that TAT-Cre translocated across the cell membrane into endometrial cells and caused excision of mT and subsequent expression of mG (Figure 2D). It is worth to mention that recombined cells were retained in the uterus for 4 months after TAT-Cre administration, indicating that this system is suitable for generation of stable Cre-recombined cells into the uterus.

### In vivo TAT-CRE mediated Pten/p53 deletion in endometrial cells leads to development of high-grade carcinomas and metastatic carcinosarcomas

Once demonstrated that TAT-Cre was able to locally promote LoxP recombination of floxed genes, we evaluated the impact of double Pten/p53 deletion, we intravaginally injected Pten^+/+^; p53^+/+^(WT)^;^ Pten^f/f^; p53^+/+^(PTENKO), Pten^+/+^; p53^f/f^ (p53KO), Pten^f/f^; p53^f/f^ (dKO) mice with TAT-Cre to induce deletion of floxed alleles. Four months after injection mice were sacrificed and submitted to pathological examination. Macroscopic observation of endometria showed an enlargement and uterine bulges in dKO mice, but no apparent macroscopic alterations in the rest of genotypes (Figure 3A). Next, endometria from different genotypes were subjected to histopathological evaluation (Figure 3B). As we previously found in tamoxifen-induced model of P53 deletion, no sign of endometrial lesions was observed in p53KO. The majority of PTENKO endometria displayed endometrial hyperplasia, EIN or low grade endometroid endometrial carcinoma (EEC1) displaying cribriform pattern to lesser extent (Figure 3C). It is worth mentioning that this results are similar to those observed after epithelial specific PTEN ablation in our tamoxifen-inducible model [23]. In contrast, dKO displayed high grade EEC (EEC3) and, 50% of mice not only had epithelial tumors, but also displayed areas with a mixed biphasic epithelial and sarcomatous cell component that were diagnosed as UCS. When analyzing the sarcomatous component of these tumors in greater detail, it was observed a high mitotic rate and a marked nuclear pleomorphism, a hallmark of this aggressive type of EC (Figure 3D).

Further pathological analysis of dKO mice revealed the presence of lung lesions in 3 out of the 9 mice injected (Figure 3E). To determine such lesions as metastatic spread of UCS, we performed an immunobiological analysis with CK8, ERα and PAX8 antibodies (Figure 3F). Positive CK8 staining evidenced the epithelial origin of metastasis. To further confirm that the lesions corresponded to endometrial metastases and to rule out the possibility of a primary lung lesion, we also carried immunohistochemistry to detect ERα and PAX8. Positive staining for these markers is indicative of endometrial metastasis [25,26].

### Uterine carcinosarcomas arise from Pten/p53 deficient endometrial epithelial cell compartment

Although all the above presented results strongly suggest that double deletion of Pten and p53 causes the development of UCS, we could not rule out the possibility that TAT-Cre ablation of Pten and p53 observed in stromal cells could lead to development of uterine sarcomas. In such scenario, biphasic tumors with mixed sarcomatous and epithelial cells could arise from double Pten/p53 deficient stromal and epithelial cells, respectively. To demonstrate that mixed tumors arise from epithelial cells that underwent EMT to originate sarcomatous component of the tumor, we performed transplantation experiments of epithelial cells of different genotypes mixed with wild type stromal cells (Figure 4A). From one side, we isolated and cultured stromal and myometrial cells from C57BLACK/6J mice (WT stroma) and, from the other side, we isolated and cultured epithelial cells from Cre:ER^T-/-^; Pten^f/f^; p53^f/f^ (WT epithelial cells), Cre:ER^T+/-^; Pten^+/+^; p53^f/f^ (P53 KO epithelial cells), Cre:ER^T+/-^; Pten^f/f^; p53^+/+^(PTEN KO epithelial cells) and Cre:ER^T+/-^; Pten^f/f^; p53^f/f^ (double PTEN/P53 KO epithelial cells) mice were injected with tamoxifen 24 hours prior isolation. We have previously demonstrated that single tamoxifen injection induces recombination specifically in epithelial cells into the uterus[23]. To ensure correct deletion of both Pten and p53, we performed a western blot analysis of epithelial cells prior transplantation (Figure 4B). Then, WT stromal cells from C57BLACK/6J mice were mixed with epithelial cells from each of the 4 genotypes and subcutaneously injected in SCID mice (Figure 4A). By using this approach, we avoided the presence of mutant stromal cells. After injection, the growth of xenotransplants was monitored weekly. Xenotransplants containing WT stroma and double Pten/p53 KO cells exhibited a dramatically increase rate of growth, reaching 1 cm^3^ size after 10 weeks (Figure 4C-4D). At this point mice required euthanasia and xenotransplants were removed. Observation of Pten/p53 dKO xenotransplant confirmed a marked increase of size over single PTENKO, P53KO or WT xenotransplants (Figure 4E). Then, dissected xenotransplants were processed for pathological evaluation. All neoplasms generated from dKO xenotransplants contained sarcomatous and carcinomatous areas. The carcinomatous component of dKO lesions was formed by a mixture of low-grade and high-grade endometrioid adenocarcinomas displaying cribriform pattern and were diagnosed as UCS (Figure 5A). In the later ones, CK8 immunostaining revealed the presence of invadopodia, indicative of its invasive phenotype (Figure 5B). Besides CK8, carcinomatous compartment was also positive for ERα or PAX8 expression (Figure 5C). In contrast sarcomatous compartment was negative for CK8, ERα or PAX8 expression and showed homologous squamous differentiation and heterologous components such as cartilaginous differentiation (Figure 5D). Both carcinomatous and sarcomatous compartment were proved to be negative for PTEN immunostaining (Figure 5E).

**Figure 4.**
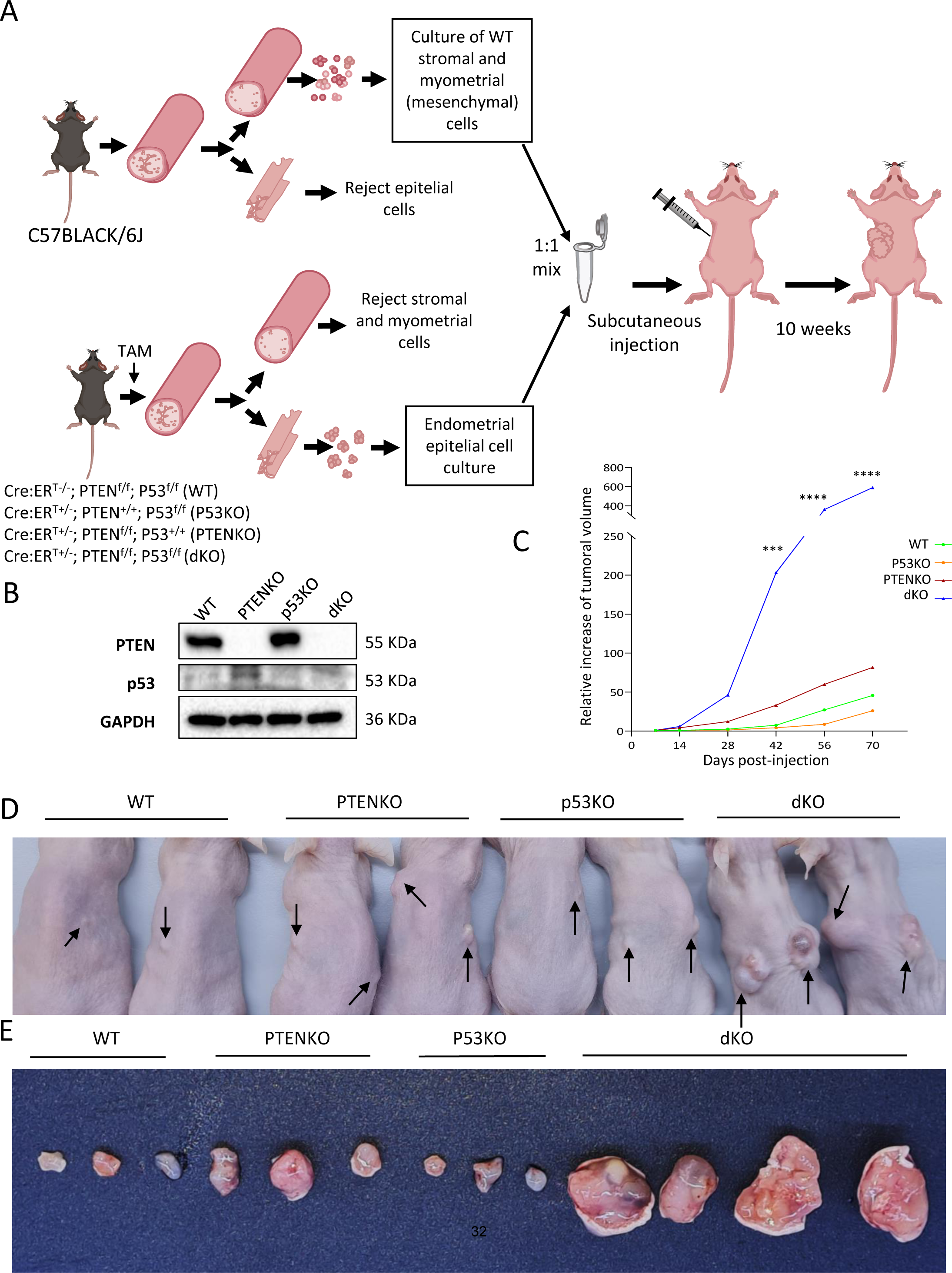
Endometrial xenotransplants contaiing wild type stromal/myometrial cells and Pten/p53 double deficient epithelial cells develop UCS. **(A)** Diagram depicting the protocol for extraction and culture of stromal and myometrial cells from the C57BLACK/6J mouse (WT) and endometrial epithelial cells from Cre:ER^T-/-^; Pten^f/f^; p53^f/f^ (WT), Cre:ERT^+/-^; P’ten^+/+^; p53^f/f^ (p50KO), Cre:ERT^+/-^; Pten^f/f^; p53^+/+^ (PTENKO) and Cre:ERT^+/-^; Pten^f/f^; p53^f/f^ (dKO). Gene deletion in vivo was induced by tamoxifen (TAM) injection. For subcutaneous injection, epithelial cells from different genotypes were mixed 1:1 with WT stromal and myometrial cells and subcutaneously injected in SCID immunosuppressed mice. **(B)** Western blot analysis of of PTEN and p53 from lysates of the epithelial cells used for the generation of xenotransplants. The GAPDH is used as loading control. **(C)** Measurement of xenotransplant growth over time. ***p-value<0.001 and ****p-value<0.0001, using a two-way ANOVA analysis, followed by a Tukey multiple comparison test. **(D)** Representative image of SCIDs mice 10 weeks after subcutaneous injection of mixed myometrial/stroma cells with epithelial cells from Cre:ER^T-/-^; Pten^f/f^; p53^f/f^ (WT), Cre:ERT^+/-^; P’ten^+/+^; p53^f/f^ (p50KO), Cre:ERT^+/-^; Pten^f/f^; p53^+/+^ (PTENKO) or Cre:ERT^+/-^; Pten^f/f^; p53^f/f^ (dKO). **(E)** Representative images of dissected xenotransplants from the 4 combinations of endometrial epithelial cells with stromal and myometrial cells. Data corresponds to the values of two independent experiments with a total of n=9 animals per condition and between 14 and 18 injected flanks per condition.

**Figure 5.**
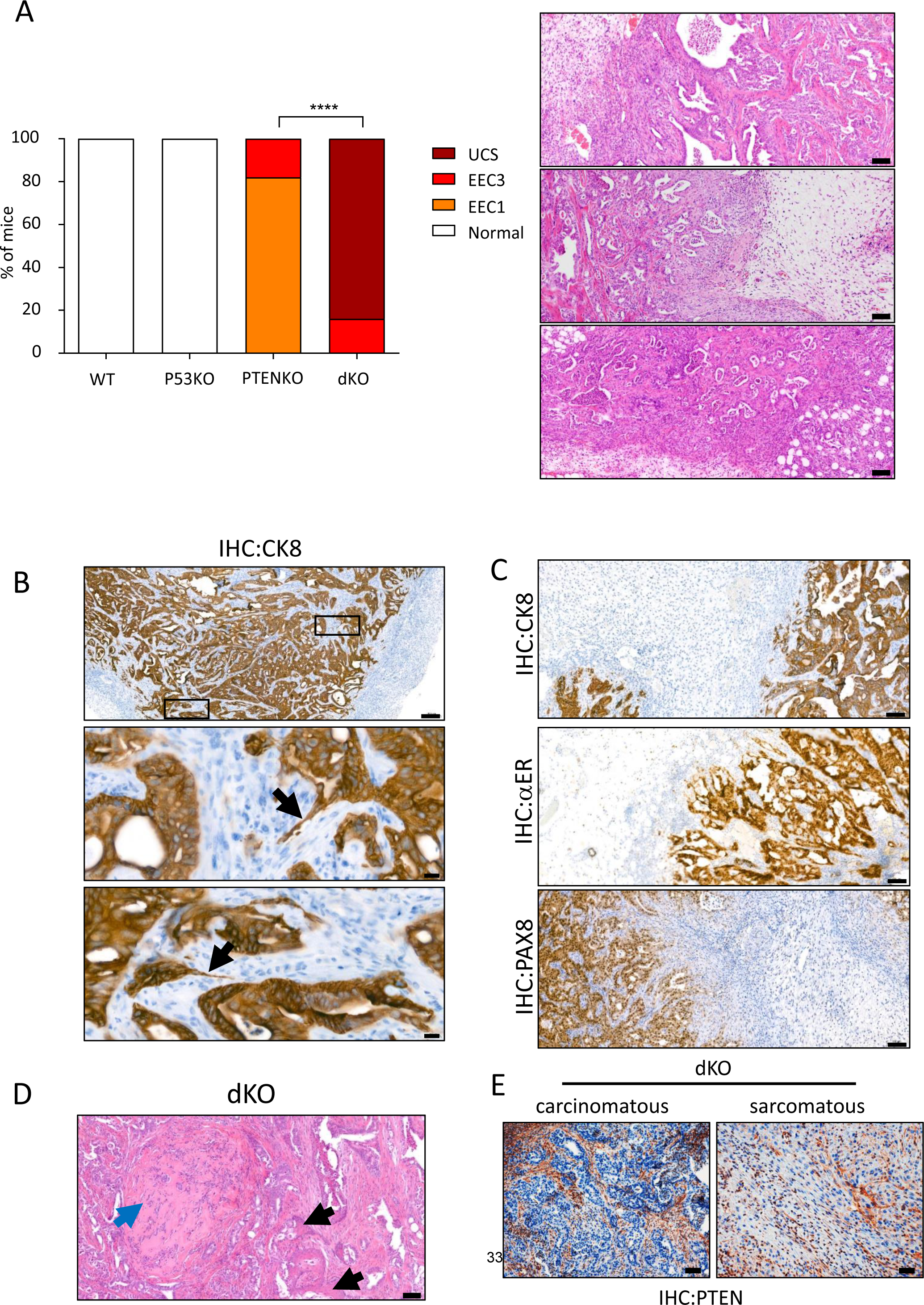
Xenografts arised from PTEN and p53 deficient endometrial epithelial show features of UCS. **(A)** Left, quantification of lesions observed in xenografts generated from Pten^+/+^; p53^+/+^ (WT), Pten^+/+^; p53^f/f^ (p53KO), Pten^f/f^; p53^+/+^ (PTENKO), and Pten^f/f^; p53^f/f^ (dKO) cells 10 weeks after transplantation. ****p<0.0001, according to the Xi^2^ test, followed by Fisher’s exact test. EIN (Endometrial Intraepithelial Neoplasia), CEE-1 (Grade 1/Low Grade Endometrioid Endometrial Carcinoma), EEC-3 (Grade 3/High Grade Endometrioid Endometrial Carcinoma), UCS (Uterine Carcinosarcoma). Right, representative hematoxylin-eosin images of 3 different subcutaneous xenografts sohwing UCS. Scale Bars: 20µm. **(B)** Representative images of CK8 immunohistochemistry of subcutaneous xenografts generated from Cre:ERT^+/-^; PTEN^f/f^; P53^f/f^ (dKO) endometrial epithelial cells showing some of the invadopodia (in black arrows). Images at 5x and 40x. Sclae bars: 200µm and 20µm. **(C)** Illustrative images of cytokeratin 8 (CK8), estrogen receptor alpha (ERα), and Paired Box 8 protein 8 (PAX8) staining of biphasic regions of carcinosarcomas developed in non-orthotopic xenografts after 10 weeks of subcutaneous injection of endometrial epithelial cells from Cre:ERT^+/-^; PTEN^f/f^; P53^f/f^ (dKO) mice. Images at 10x. Scale bars: 100µm. **(D)** Representative image showing squamous differentiation of the endometrial epithelial cells (black arrows) and sarcomatous regions with heterologous (cartilaginous) differentiation (blue arrow). Images at 10x. Scale bar: 100 µm. **(E)** Representative images PTEN immunohistochemistry in the sarcomatous component (right) and carcinomatous component (left) of UCS generated from Cre:ERT^+/-^; PTEN^f/f^; P53^f/f^ (dKO) endometrial epithelial cells. The positive labeling of the wild-type mesenchymal cells that were introduced mixed with the epithelial cells when generating the subcutaneous tissues can be seen. Images at 10x. Scale bars: 100µm.

Likewise conditional KO endometria treated with TAT-Cre, tumors arised from PTENKO xenotransplants were restricted to low-grade endometroid endometrial carcinomas and P53KO did not exhibit any apparent lesion (Supplementary Figure 4).

### Pten/p53 deficient UCS xenotransplants display high proliferation rates, nuclear pleomorphism, and metastatic potential

As aforementioned, high proliferation rates and nuclear pleomorphism are associated with aggressive phenotype of high-grade endometrial adenocarcinomas and UCS. To demonstrate these two features in UCS raised from xenotransplants, we performed Ki-67 immunostaining and analysis of nuclear area. Ki-67 immunostaining demonstrated a significant increase of double Pten/p53 tumors over those lacking only Pten (Figure 6A). Measurement of nuclear areas of double Pten/p53 evidenced a marked increase in nuclear pleomorphism when compared with tumors deficient for Pten (Figure 6B).

**Figure 6.**
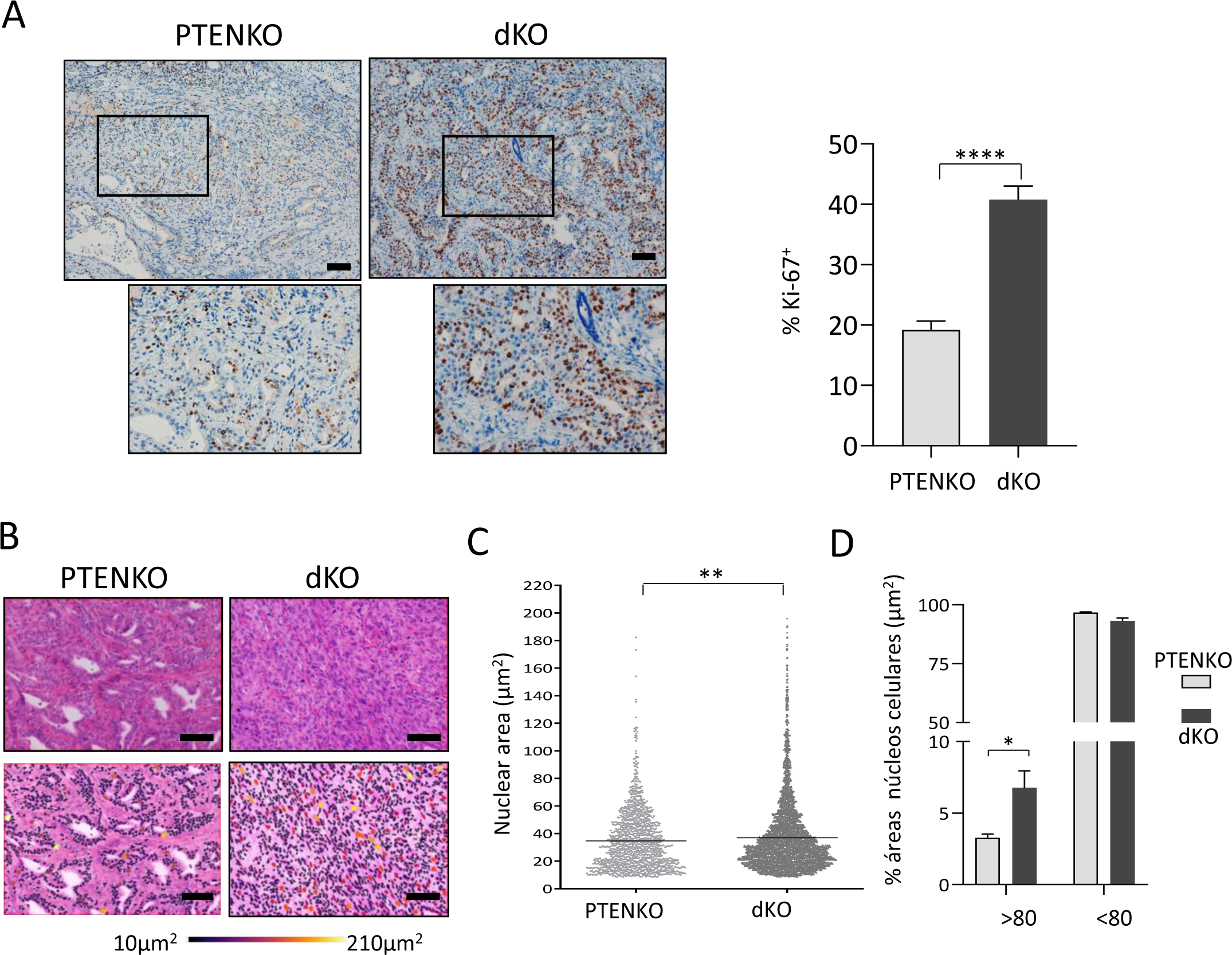
Tumors deficient for Pten and p53 have a higher proliferation rate and nuclear pleomorphism than tumors lacking PTEN. **(A)** Representative images and quantification of Ki-67 staining xenografts after 10 weeks of subcutaneous injection of endometrial epithelial cells from Cre:ERT^+/-^ mice; Pten^f/f^; p53^+/+^ (PTENKO) and Cre:ERT+/-; Pten^f/f^; p53^f/f^ (dKO). Images at 10x. Scale bars: 100µm bars. Data from n=3 xenograft for each group. Values are expressed as the mean ± s.e.m. ****p<p0.0001, based on t-test analysis. **(B)** Upper images are representative images of hematoxylin-eosin staining of subcutaneous tumors generated from Cre:ERT^+/-^ endometrial epithelial cells; Pten^f/f^; p53^+/+^ (PTENKO) and Cre:ERT^+/-^; Pten^f/f^; P53^f/f^ (dKO). Lower pannel are images analyzed with the QuPath software, where the nuclei are marked with a color gradient from the largest (210 µm2) to the smallest (10 µm2) nuclear area size. Scale bars: 100µm. **(C)** Measurements of the nuclear areas of the subcutaneous endometrial tumors in µm^2^. The data was obtained from 3 different regions counts in independent tumors using the QuPath software. Error bars represent mean ± s.e.m. **p<0.01, using an unpaired t-test followed by a Bonferroni multiple comparison. **(D)** Quantification of the percentage of nuclear areas larger and smaller than 80µm2 in PTENKO and dKO tumors. Error bars represent mean ± s.e.m *p<0.05, according to two-way ANOVA followed by Sidak multiple comparison. test

### In vivo CRISPR/Cas9 disruption of Pten and p53 in endometrial epithelial cells results in the development of metastatic UCS

We have recently demonstrated that in vivo electroporation of CRISPR/Cas9 ribonucleoproteins is a suitable tool for epithelial-cell specific disruption tumor suppressor genes in the uterus [17]. By applying this methodology, we sought to investigate whether concurrent disruption of Pten and p53 in epithelial cells of wild type mice uteri would lead to development of UCS. Three wild type C57BACK6/6J mice were co-electroporated with Pten- and p53-targeting RNPs and 4 months later mice were sacrificed and examined for the presence of EC and metastatic dissemination. All three electroporated mice developed metastatic EC. Histopathological evaluation revealed revealed the presence of high-grade tumors displaying carcinomatous and sarcomatous components in all 3 electroporated mice and were diagnosed of UCS (Figure 7A). To confirm the presence of Pten and p53 edits causing the disruption of the two targeted tumor suppressor genes, DNA extracted from one primary tumor (located in the uterus) and one metastatic outgrowth was submitted to Amplicon Next Generation Sequencing (NGS) (Figure 7B). In both primary and metastatic samples, CRISPR/Cas9 editing resulted in a high number of insertions and deletions (indels) in Pten and p53 tumour suppressor genes. However, a significant percentage of reads corresponding to wild type, non-edited Pten and p53 sequences were also detected. Consistently, Pten and p53 immunohistochemistry on consecutive tissue sections demonstrated a simultaneous loss of Pten and p53 expression in both primary and metastatic parenchymal cancerous cells co-existing with wild-type cells retaining Pten or p53 expression corresponding to stromal, endothelial, or inflammatory cells (Figure 7C).

**Figure 7.**
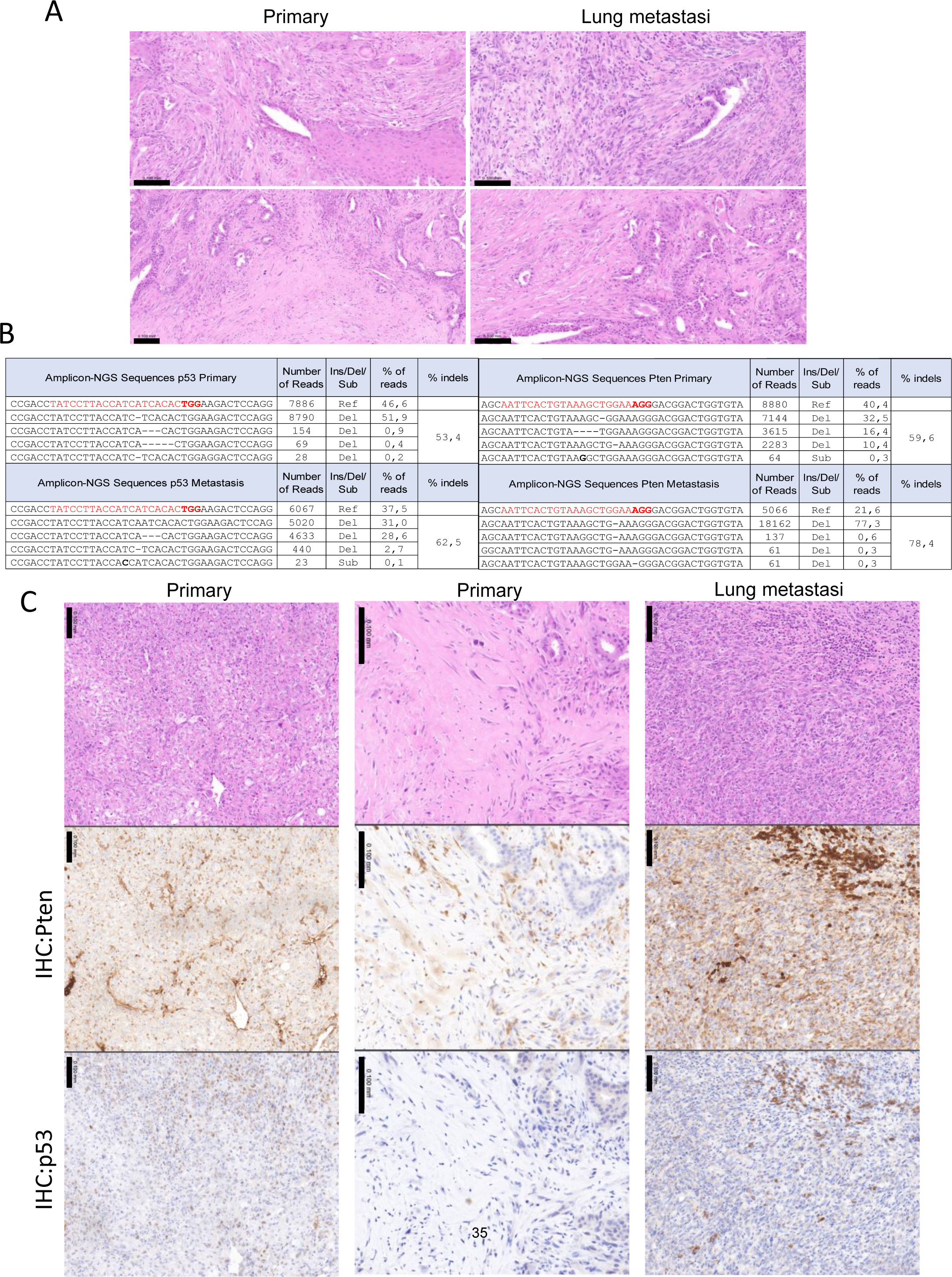
In vivo CRISPR/Cas9 editing of Pten and p53 results in UCS development. **(A)** Representative hematoxylin-eosin staining on sections corresponding to two primary tumors and two lung metastatic outgrowths dissected from mice co-electroporated with Pten and p53 targeting RNPs. **(B)** Summary of Pten and p53 amplicon-NGS results showing the sequence, number of reads, type of edits and percentage of edits compared to the wild type (Ref) sequence obtained from sequencing amplicons from a primary tumor and a metastatic outgrowth. **(C)** Representative hematoxylin-eosin staining, Pten and p53 immunohistochemistry on consecutive sections corresponding to two primary tumors and one lung metastatic outgrowth.

## DISCUSSION

As most malignancies, EC arises from the accumulation of driver mutations that confer the hallmarks of cancer cell phenotype [27]. Whole genome sequencing have unveiled or confirmed previously known mutations in EC[6]. However, functional validation of the identified alterations as a driver mutation itself or in combination with other alterations is still a challenge in cancer research. GEMM play a pivotal role in the understanding of candidate driver cancer gene function. Many studies have reported conditional murine models for individual deletion of Pten or p53 or combined deletion with other frequently altered genes in EC [7]. However, most of these models present a series of limitations, which have provided heterogeneous and, sometimes, contradictory results. For instance, depending on the Cre recombinase expressing mice used for conditional ablation of Pten, some models develop invasive endometrial adenocarcinomas [16], in situ endometrial adenocarcinomas [23] or endometrial hyperplasia [28]. Disparities have been also reported for p53 mouse models. To date, there is only one mouse model addressing the effects of individual p53 deficiency in EC development[15]. Mice with endometrium-specific deletion of p53 developed carcinomas representing all type II histological subtypes, including serous, clear cell, undifferentiated EEC and UCS[15]. However, other studies in which p53 has been deleted along with other tumor suppressor genes have also shown that loss of p53 alone is not sufficient to cause EC. Conditional deletion of Pten/p53[29], Cdh1/p53[30], Pot1a/p53[31], or Rb1/p53[32] in mouse endometrium resulted in EC but, in all these studies, control mice lacking only p53 had no signs or low incidence and penetrance of EC. More recently, another study demonstrated that conditional loss of p53 in the presence of PIK3CA^H1047R^ mutation drives hyperplasia and endometrial intraepithelial carcinoma, but control mice lacking only p53 had no endometrial lesions[29]. In line with these reports, using TAT-Cre mediated deletion of p53, we have also found that loss of p53 is not sufficient to cause EC. Although we and others have found that ablation of p53 is not enough to cause EC, human endometrial neoplasia harboring p53 mutations exhibit the most aggressive forms of EC with the worst prognostic. Obviously, carcinogenesis is a multistep process in which other molecular alterations will join p53 to cause EC. Histologically, human p53-mutated tumors are high-grade EECs, serous carcinomas or UCS. To date, there is not existing mouse model in which specific deletion of Pten and p53 in endometrial epithelial cells has been reported. The only existing mouse model addressing the effects p53 and Pten[16] deletion mice reported the presence invasive EECs, with no signs of UCS, clear cell or serous carcinomas. In contrast, we demonstrate that TAT-Cre mediated Pten/p53 ablation or CRISPR/Cas9-mediated editing results in a high percentage of high-grade EECs and UCS, thereby recapitulating histopathology of human tumors. Regarding UCS, TAT-Cre and CRISPR/Cas9 systems provided new insight to this histologic manifestation of ECs. First, intrauterine delivery of TAT-Cre provided the first evidence that, among all combinations of p53 deficiency with other tumor suppressors genes deficiency, simultaneous Pten and p53 deletion is enough to cause this type of malignancy. Second, RNA sequencing demonstrated that double Pten/p53 deletion is enough to trigger molecular changes associated to EMT. Such changes are consistently translated to the development of UCS in which epithelial cells are transformed to mesenchymal cells, thereby originating the sarcomatous component of UCS. Accordingly, xenografting of both double Pten/p53 deficient epithelial cells with wild type stroma cells and epithelial-cell specific editing of Pten and p53 demonstrate the epithelial origin of both carcinomatous and sarcomatous compartments of UCS.

The discrepancies in EC development among mouse models with conditional deletion of Pten, p53 or Pten/p53 can be explained by different spatial and temporal activity of Cre recombinase used in each study. One key advantage of TAT-Cre administration and CRISPR/Cas9 technology is the temporal control of tumor suppressor gene deletion. There is a still limited amount of Cre expressing mouse that allow both specific and inducible control of floxed gene recombination in the uterus. For instance, Cre expression driven by progesterone receptor (*Pgr)*[33], anti-Müllerian hormone type 2 receptor *(Amhr2)*[34(p1)], and *Wnt7a* promoters display activity before or just after birth. Others such as the *Spfrr2*[35]*, BAC-Sprr2f*[36] or lactoferrin (*Ltf*) promoter driven Cre expression[37] is activated with the beginning of estrous cycle which may also be a premature age for human EC modelling. By simply choosing the age of mice in which TAT-Cre is intravaginally injected or CRISPR/Cas9 RNPs are electroporated, deletion of genes of interests can be achieved at different points of mouse lifetime. Since the incidence of cancers dramatically increases with age, we think these techniques provide a more suitable method for EC recapitulation. So far, our previously reported tamoxifen-inducible[23] and a tetracicline-inducible[38],[32] mouse model are the only two reported systems for inducible Cre activity. Unfortunately, most of them have other limitations that limit their use to model EC. Although inducible, in these existing models Cre activity ablation LoxP-flanked alleles is not restricted to the uterus, which difficult the study of tumor suppressor gene deletion in EC.

Regarding spatial control of Cre activity, to date, there is not existing mouse model allowing specific deletion of LoxP-flanked genes in epithelial cells, without affecting stromal/myometrial into the uterus or in other extrauterine tissues[7]. Inactivation of LoxP-flanked tumor suppressor genes specifically in the epithelial component is crucial to understand the autonomous mechanisms involved in EC. As in other types of neoplams, deletion of tumor suppressor genes specifically in epithelial compartment or in epithelial and stromal/myometrial cells of the uterus may have different consequences. For instance, it has been demonstrated that epithelial specific deletion of Pten using Ltf-driven Cre expression a causes endometrial hyperplasia whereas Pten ablation in stromal and epithelial cells using the Pgr-driven Cre expression[16] leads to invasive endometrial carcinomas[28]. These results strongly suggest that Pten deletion in endometrial stromal cells is likely to affect epithelial cell tumor progression.

As most cancers, EC arise from accumulation of mutations in a single cell. Initiated cell and/or its daughter cells progressively acquire the features of cancer phenotype. Most of available Cre expressing strains used to model EC cause a massive ablation of LoxP-flanked genes in all Cre expressing cells into the uterus (either specifically in the epithelial compartment or in both stromal, epithelial, or myometrial cells). In contrast, TAT-Cre recombination of floxed genes or CRISPR/Cas9 editing also renders a cell mosaicism in which non-recombined wild type cells co-exist side-by-side with recombined cells. Even mouse models in which Cre recombinase is restricted to epithelial cells such as the Ltf-Cre or the Spfrr2-Cre expressing mice, Cre induces LoxP-flanked genes deletion in virtually all epithelial cells, leaving no Pten proficient epithelial cells. Considering that loss of Pten in all stromal cells may affect EC development[28], is easy to speculate the complete absence of Pten wild type epithelial cells may also affect EC development and progression. Here, we have demonstrated that TAT-Cre intravaginal administration results in mosaic Pten deletion in epithelial and stromal cells, leaving Pten expression intact most of both cells of both compartments. Likewise, epithelial cell specific editing of p53 and Pten by the CRISPR/Cas9 system, generates a mosaicism in which low amount or edited cells co-exist with wild type, non-edited cells.

Besides all the above mentioned Cre-expressing mouse models, the only reported method for local delivery of Cre recombinase is mediated by intrauterine injection of Adenovirus carrying Cre transgene (Ad-Cre)[39,40]. However, a significant limitation of this Ad-Cre approach was an extremely low efficiency. Semi-quantitative PCR analysis indicated that the efficiency of recombination was <1% in endometrial epithelial cells, with no recombination detectable in ∼50% of animals. In contrast, 100% of mouse in which TAT-Cre was injected displayed recombined cells.

Collectively, our results provide the first functional demonstration that simultaneous Pten and p53 mutation is sufficient to trigger EMT that is consistently translated in the development UCS, the EC histological subtype of EC in which this process systematically is observed.

## ACKNOWLEDGEMENTS.

Supported by grants and PID2019-104734RB-I00 from Spanish Ministerio de Ciencia, Innovación y Universidades,

## AUTHOR CONTRIBUTION

Conceptualization, RN and X.D.; methodology, RN, GA, ARM, MVS, APG, AY, and X.D.; investigation, SG, RN, ARM, MVS; validation, all authors; formal analysis, SG, GA, AY, JE, ME, XMG, XD; resources, XMG, XD.; writing, RN, JC, XD.; supervision, XD.; main project administration, XD.; funding acquisition, XD.

