## Supplementary Figures for "*In vivo* intra-uterine delivery of TAT-fused Cre recombinase and CRISPR/Cas9 editing unveil histopathology of Pten/p53-deficient endometrial cancers"

### Supplementary Figure 1

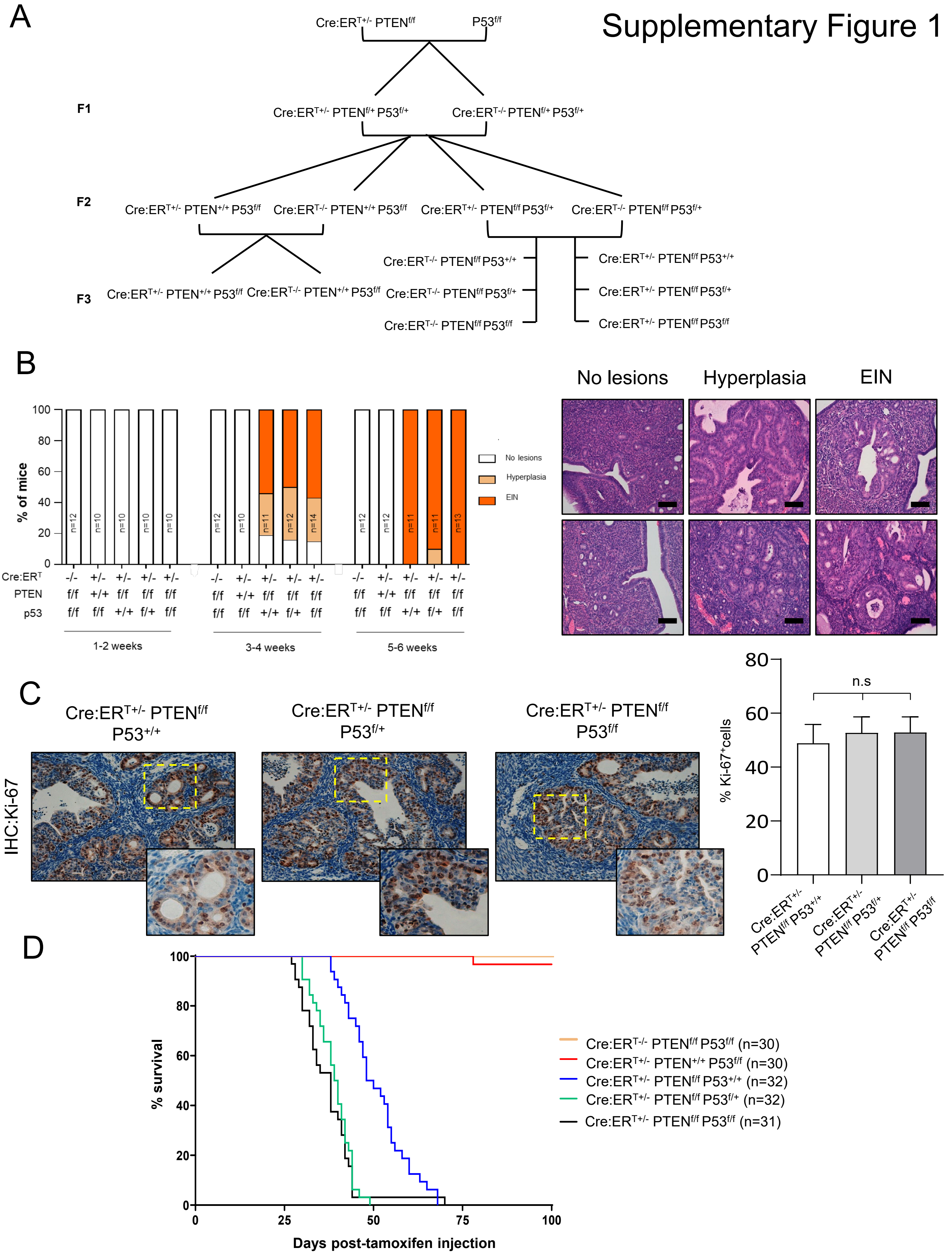

**Supplementary Figure 1. In vivo Tamoxifen-inducible deletion of PTEN and P53 leads to the development of hyperplasia and non-invasive intraepithelial neoplasia. (A)** Breeding protocol for the generation of murine models Cre:ERT+/-; PTENf/f; P53f/f (dKO), Cre:ERT+/-; PTENf/f; P53+/+ (PTENKO) and Cre:ERT+/-; PTEN+/+; P53f/f (p53KO). **(B)** Quantification and representative hematoxylin-eosin images of endometrial lesions for the indicated groups of mice. ****p<0.0001, according to the $Xi^2$ test, followed by Fisher's exact test. EIN (Endometrial Intraepithelial Neoplasia). **(C)** Representative Ki-67 immunohistochemistry images of $Pten^{+/+}$; Cre:$ER^{T+/-}$ $Pten^{+/+}$ $p53^{f/f}$, Cre:$ER^{T+/-}$ $Pten^{f/f}$ $p53^{+/+}$, and Cre:$ER^{T+/-}$ $Pten^{f/f}$ $p53^{f/f}$ uterine sections 6 weeks after tamoxifen injection. **(D)** Kaplan-Meier plot showing survival of animals Cre:$ERT^{-/-}$; $PTEN^{f/f}$; $P53^{f/f}$ (WT), Cre:$ERT^{+/-}$; $PTEN^{+/+}$; $P53^{f/f}$(P53KO), Cre:$ERT^{+/-}$; $PTEN^{f/f}$; $P53^{+/+}$ (PTENKO), Cre:$ERT^{+/-}$; $PTEN^{f/f}$; $P53^{f/+}$ and Cre:$ERT^{+/-}$; $PTEN^{f/f}$; P53/f(dKO). p-value <0.01. Statistics performed with the Log-rank test (Mantel-Cox).

### Supplementary Figure 2

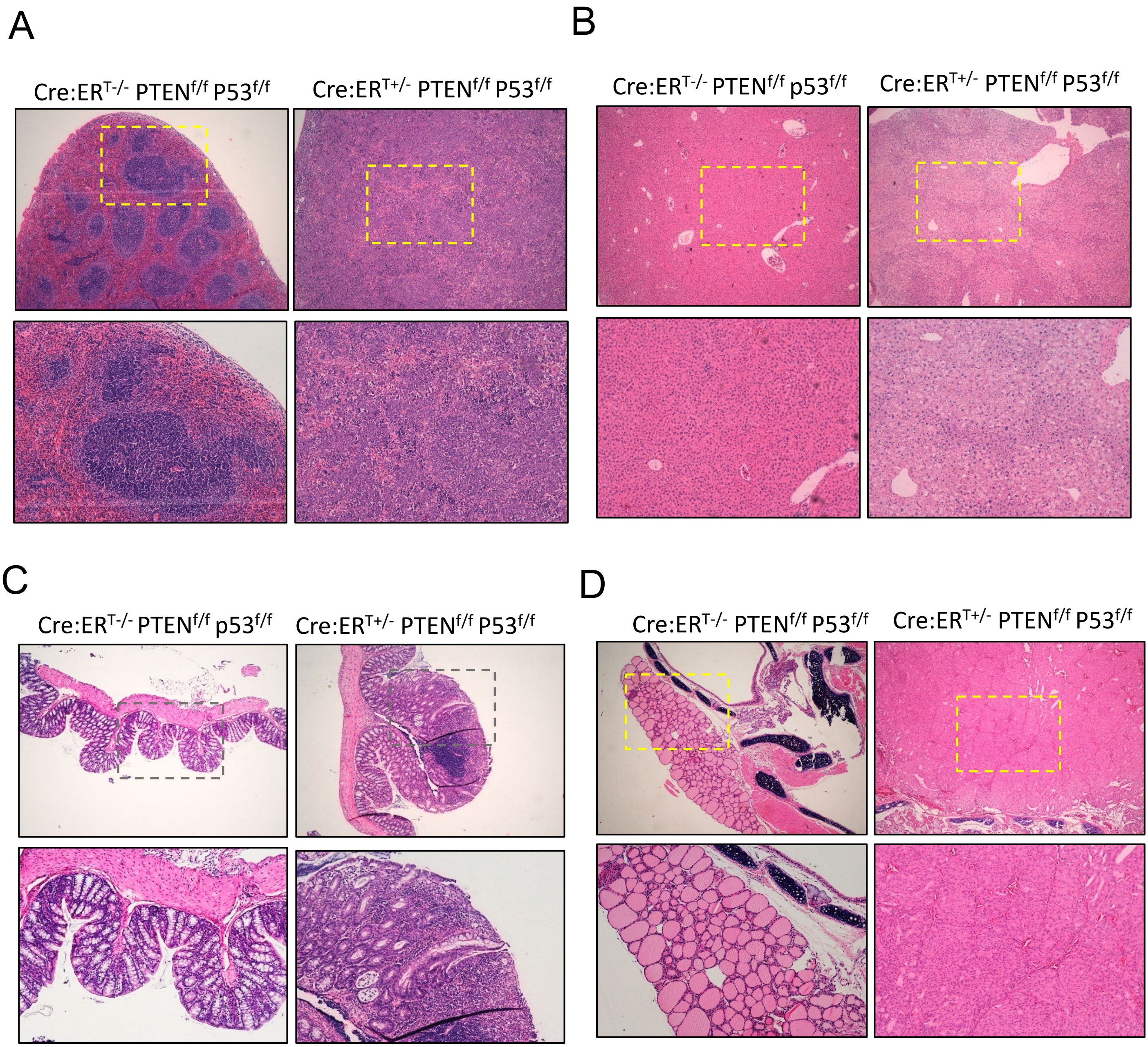


**Supplementary Figure 2. Histopathological study of lesions observed in double Pten/p53 knock-out mice**. Representative images of lymphoma **(A)**, hepatocellular displasia **(B)**, colon adenomas **(C)**, or severe thyroid hyperplasia **(D).**

### Supplementary Figure 3

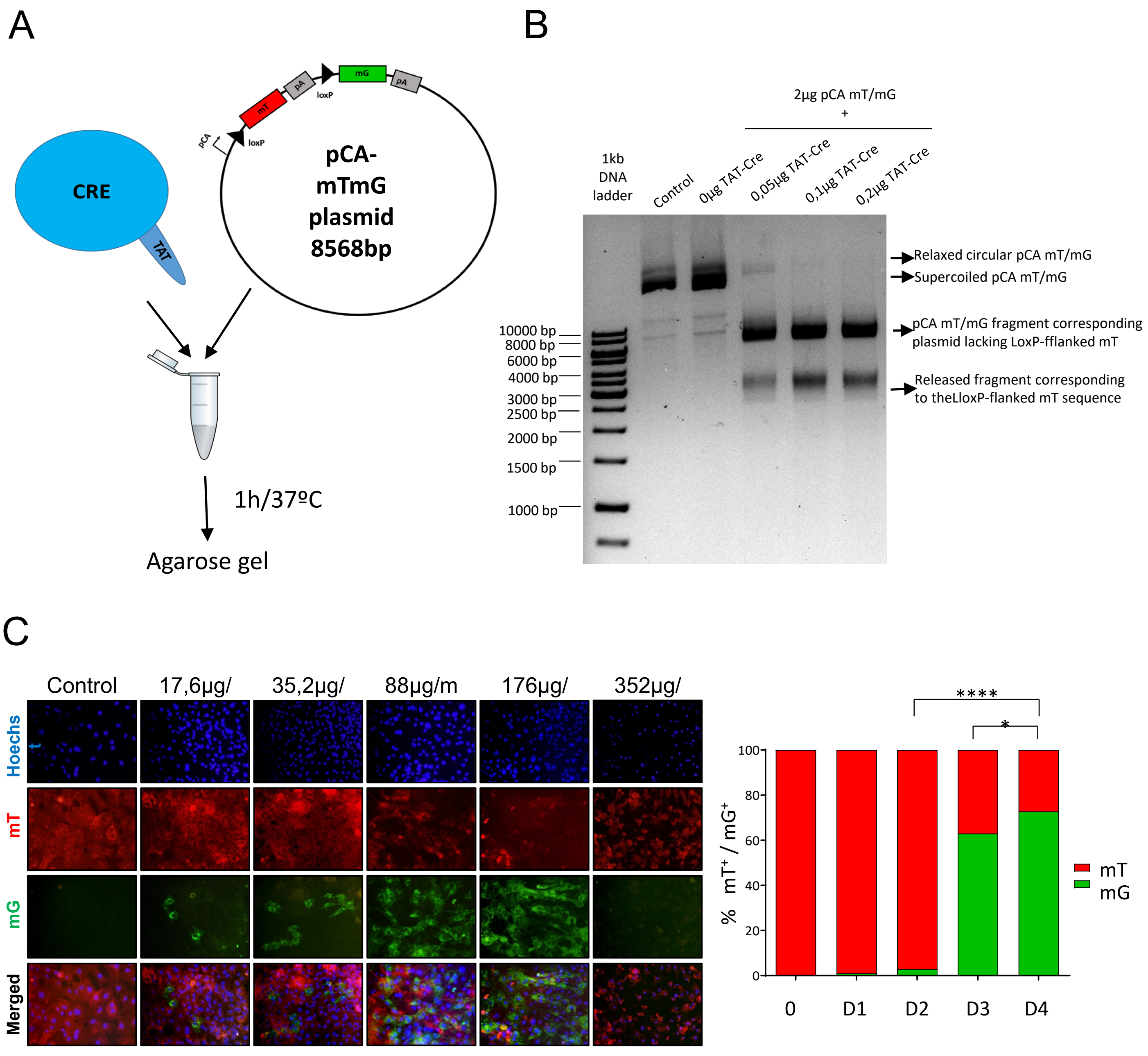


**Supplementary Figure 3. TAT-Cre exhibits recombinase activity in vitro. (A)** Scheme of the in vitro assay for the analysis of TAT-Cre recombinase activity using the pCA-mTmG plasmid. **(B)** Representative image of an agarose gel showing the DNA fragments of pCA-mT/mG plasmid generated by TAT-Cre activity. **(C)** Representative images and quantification of tdTomato positive ($mT^+$) or GFP positive ($mG^+$) fibroblasts isolated from mT/mG$^{f/f}$ mouse ears 4 days after being treated with the following TAT-CRE concentrations: 0µg/mL (0), 17.6µg/mL (D1), 35.2µg/mL (D2), 88µg/mL (D3), 176µg/mL (D4) and 352µg/mL (D5). Images captures at 20X. Data from n=3 independent experiments. *$p<0.5$ ****$p<0.0001$, using a one-way ANOVA analysis, followed by a Bonferroni multiple comparison test.

### Supplementary Figure 4

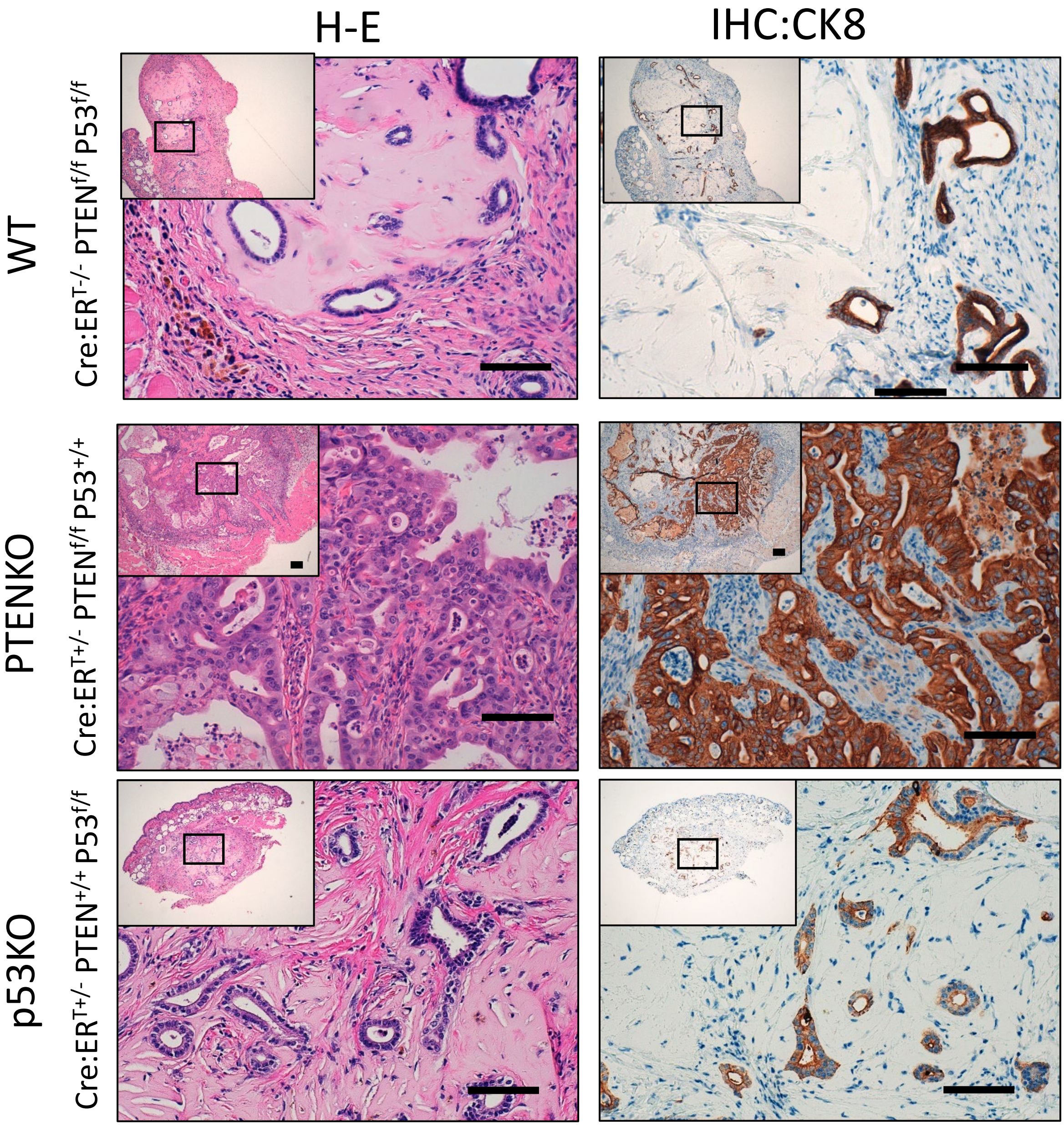


**Supplementary Figure 4. Histopathology of WT, PTENKO or p53KO xenotransplants.**. Representative images of hematoxylin-eosin images (H-E) and cytokeratime-8 (CK8) immunohistochemistry on endometrial tissues and lesions developed from xenotransplated endometrial epitelial cells of the indicated genotypes.
