## Supplementary methods for "*In vivo* intra-uterine delivery of TAT-fused Cre recombinase and CRISPR/Cas9 editing unveil histopathology of Pten/p53-deficient endometrial cancers"

**Table SM1.** Genotyping primers and PCR conditions.

| Strain | Primers | | PCR protocol | | | Resulting bands | |
|---|---|---|---|---|---|---|---|
| | | | T (ºC) | times | Cycle | Genotype | bsands |
| Cre:ER$^{T}$ | Fwd<br>Rev | ACG AAC CTG GTC GAA ATC GT GCG<br>CGG TCG ATG CAA CGA GTG ATG AG | 94 ºC | 2’ | 1 | Cre:ER$^{T-/-}$<br>Cre:ER$^{T+/-}$ | no band<br>350 bp |
| | | | 94 ºC | 45’’ | 32 | | |
| | | | 65 ºC | 45’’ | | | |
| | | | 72 ºC | 45’’ | | | |
| | | | 72 ºC | 5’ | 1 | | |
| *Pten* floxed | Fwd<br>Rev | CAA GCA CTC TGC GAA CTG AG<br>AAG TTT TTG AAG GCA AGA TGC | 94 ºC | 3’ | 1 | PTEN$^{+/+}$<br>PTEN$^{f/+}$<br>PTEN$^{f/f}$ | 156 bp<br>156 y 328 bp<br>328 bp |
| | | | 94 ºC | 30’’ | 35 | | |
| | | | 60 ºC | 1’ | | | |
| | | | 72 ºC | 2’ | | | |
| | | | 72 ºC | 2’ | 1 | | |
| *p53 floxed* | WT<br>Rev | CAC AAA AAC AGG TTA AAC CCA G<br>AGC ACA TAG GAG GCA GAG AC | 94 ºC | 3’ | 1 | P53$^{+/+}$<br>P53$^{f/+}$<br>P53$^{f/f}$ | 288 bp<br>288 y 370 bp<br>370 bp |
| | | | 94 ºC | 30’’ | 35 | | |
| | | | 59 ºC | 1’ | | | |
| | | | 72 ºC | 1’ | | | |
| | | | 72 ºC | 3’ | 1 | | |
| *mT/mG* | Común<br>WT<br>Mutante | CTC TGC TGC CTC CTG GCT TCT<br>CGA GGC GGA TCA CAA GCA ATA<br>TCA ATG GGC GGG GGT CGT T | 94 ºC | 2’ | 1 | mT/mG$^{+/+}$<br>mT/mG$^{f/+}$<br>mT/mG$^{f/f}$ | 330 bp<br>250y 330 bp<br>250 bp |
| | | | 94 ºC | 30’’ | 35 | | |
| | | | 57 ºC | 1’ | | | |
| | | | 72 ºC | 1’ | | | |
| | | | 72 ºC | 2’ | 1 | | |

**Table SM2.** Antibodies used for immunohistochemistry.

| Antibody | Dilution | Commertial Company | Catlao¡log | EnVision™ FLEX | Secondary antibody |
|---|---|---|---|---|---|
| **PTEN** | 1:100 | Dako | M3627 | *High pH* | EV FLEX Kit |
| **Ki-67** | 1:50 | Dako | M7249 | *Low pH* | EV FLEX Kit |
| **ERG** | RTU | Dako | IR659 | *High pH* | EV FLEX Kit |
| **αER (1D5)** | RTU | Dako | IR657 | *High pH* | EV FLEX Kit |

| | | | | | |
|---|---|---|---|---|---|
| **E-cadherin** | RTU | Dako | IR059 | *High pH* | EV FLEX Kit |
| **TTF1** | RTU | Dako | M3575 | *High pH* | EV FLEX Kit |
| **Cytoqueratin 8** | 1:200 | DSHB | AB531826 | *High pH* | Rata anti-biotin |
| **αSMA** | RTU | Dako | IR611 | *High pH* | EV FLEX Kit |
| **h-caldesmon** | RTU | Dako | GA054 | *High pH* | EV FLEX Kit |
| **Calretinin** | RTU | Dako | IR627 | *High pH* | EV FLEX Kit |
| **CD10** | RTU | Dako | GA648 | *High pH* | EV FLEX Kit |
| **Desmin** | RTU | Dako | IR606 | *High pH* | EV FLEX Kit |
| **PAX8** | 1:100 | GENOVA | AP10903 | *High pH* | EV FLEX Kit |
| **GFP** | 1:100 | Rockland | 600-101-215 | *High pH* | Cabra anti-biotina |
| **p-AKT (Ser473)** | 1:50 | Cell signalling | 3787 | *High pH* | Rabbit anti-biotin |
| **EnVision FLEX detection kit** | RTU | Dako | K8002 | - | - |
| **GOat anti-biotin** | 1:200 | Santacruz | SC-2489 | - | - |
| **Rat anti-biotin** | 1:200 | ABCAM | AB6733 | - | - |
| **Rabbit anti-biotina** | 1:200 | Jackson | 111-065-144 | - | - |
| **Estreptavidin-HRP** | 1:400 | Dako | P0397 | - | - |

**Table SM3.** Antibodies used in immunofluorescence.

| Antigen | Dilution | Commertial company | Catalog |
|---|---|---|---|
| **Phalloidin** | 1:1000 | Sigma-Aldrich | P1951 |
| **Vimentin** | 1:200 | BD bioscience | 550513 |
| **Cytoqueratin** | 1:200 | Abcam | 9377 |
| **Anti-mouse IgG Alexa Fluor™ 488** | 1:250 | ThermoFisher | A11029 |
| **Anti-mouse IgG Alexa Fluor™ 546** | 1:250 | ThermoFisher | A11010 |

**Table SM4.** Antibodies used for western blot analysis.

| Antígen | Dilution | Commertial Company | Catalog |
|---|---|---|---|
| **PTEN** | 1:1000 | Cell Signaling technology | 9188 |
| **P53** | 1:1000 | Leica | VP-P956 |
| **β-Actin** | 1:5.000 | Santa Cruz Biotechnology | sc-1616 |
| **GAPDH** | 1:20.000 | Abcam | 8245 |
| **HIS-TAG** | 1:1000 | Cell Signaling technology | 2365 |
| **Rabbit IgG-HRP** | 1:10.000 | Jackson | 111-035-003 |
| **Rabbit IgG-HRP** | 1:10.000 | Jackson | 115-035-003 |

**Table SM5.** Primers and PCR conditions for Amplicon-NGS sequencing. DNA fragments flanking the target sequence of the RNPs targeting the indicated genes were amplified with the indicated primers and PCR conditions in 50µl PCR reactions using Taq polymserase (Biotools).

| Targeted gene | | Primer | PCR Protocol | | |
|---|---|---|---|---|---|
| | | | **T (ºC)** | **Time** | **Cycles** |
| ***Pten*** | Fwd<br>Rev | TTATCTTTTTACCACAGTTGCAC<br>GTGGTTGTATCCACTTAGTGTA | 95 ºC | 2’ | 1 |
| | | | 95 ºC | 30’’ | 45 |
| | | | 55 ºC | 30’’ | |
| | | | 72 ºC | 1’20’’ | |
| | | | 72 ºC | 7’ | 1 |
| ***p53*** | Fwd<br>Rev | CCATGCTAAGCAAGTGTTGG<br>CCCTAAGCCCAAGAGGAAAC | 95 ºC | 2’ | 1 |
| | | | 95 ºC | 30’’ | 45 |
| | | | 55 ºC | 30’’ | |
| | | | 72 ºC | 1’20’’ | |
| | | | 72 ºC | 7’ | 1 |

**Figure SM1.** Images of the Coomassie blue stained acrylamide gel showing Purified TAT-Cre band. F-T; Amicon® (flow-through) eluted volume, LB OEI Medium; Lysogeny Broth Overnight Express™ Instant medium. kDa; kilodaltons. MW: BenchMark™ protein marker.

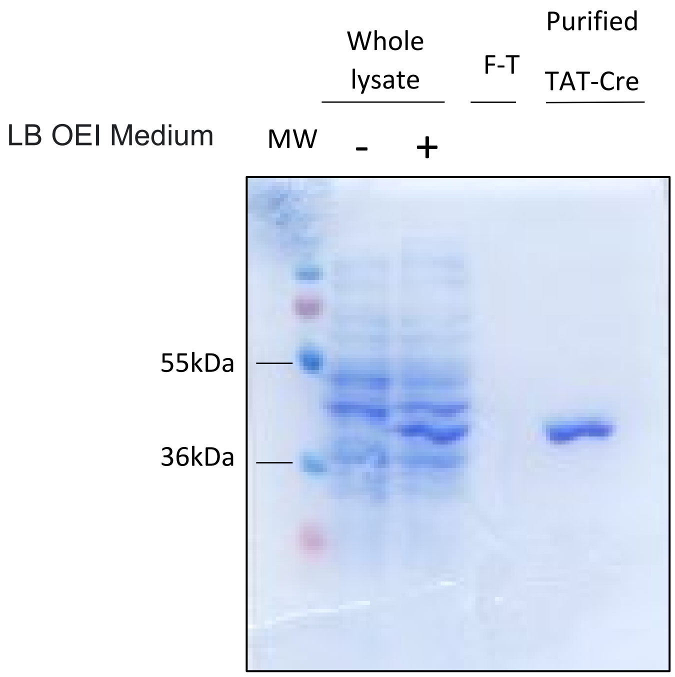
